# Molecular Dynamics Exploration of Two Full-Length IgG Antibodies: Impact of Glycosylation

**DOI:** 10.64898/2026.04.14.718417

**Authors:** Ravy Leon Foun Lin, Adam Bellaiche, Julien Diharce, Catherine Etchebest

## Abstract

Monoclonal antibodies are among the most important biomolecules in the pharmaceutic field. They could undergo several post-translational modifications, most notably N-glycosylation, yet the effect of these glycans remains poorly understood at the atomistic level, and the few existing studies focus on the prevalent immunoglobulin G1. We compared N-glycosylation in two structurally divergent monoclonal antibodies, Mab231, a murine IgG2a, and pembrolizumab, a human IgG4, which differ in species, sequence, architecture, and in the location and composition of their glycans. Using molecular dynamics simulations, we studied both antibodies with and without their glycans. In both, the glycan interactions extend beyond the Fc to reach Fab residues, more so for the more mobile Fc glycan and differently in the two antibodies. Allosteric network calculations reveal a potential impact of the glycan that can affect the Fab framework regions, which could in turn affect antigen binding. The glycans do not drastically alter the conformational landscape, and neither the inter-domain correlated motions nor the orientation of the Fab arms relative to the Fc can be resolved as glycan effects at three replicates per system. We also find that the effect of the Fc glycans on CH2 opening depends on the geometric criterion used, which underscores the need to consider full-length structures and the diversity of IgG scaffolds in glyco-engineering.

## INTRODUCTION

Monoclonal antibodies (mAbs) are proteins belonging to the immunoglobulin G (IgG) class^1^. They have become central tools in the pharmaceutical^2–4^ and biotechnology^5,6^ industries, where they are used to treat cancer^2,6,7^ and leukaemia^8,9^, and to power routine diagnostic assays such as the ELISA test^10^. Their therapeutic value stems from a single structural feature: their complementarity-determining regions (CDRs) can be tailored to recognise a unique target^5^, which minimises undesired immune responses and enables the specific detection or neutralisation of pathogens. In practice, however, some mAbs display polyspecificity, cross-reactivity, limited stability, or low binding affinity, properties that compromise the intended immune response. Designing antibodies with improved specificity, higher affinity, and greater stability therefore remains a central objective for biotechnology and pharmaceutical companies^1^. To this end, a broad set of *in vitro*, *in silico*, and *in vivo* strategies has been developed, most of them centered on the optimization of the CDRs^11–13^. Beyond antigen recognition itself, such approaches can also be used to stabilize the mAb scaffold or to improve binding specificity in complex molecular environments^1,8,10,14^.

Although the functional properties of IgGs are now well characterized, the dynamics of the full-length structure remains comparatively poorly understood^15^. Structurally, an IgG can be described as three main fragments: the crystallisable fragment (Fc) and two antigen-binding fragments (Fabs)^16–18^ (Fig. 1). Each Fab is composed of a light chain and a heavy chain, connected by disulphide bonds^17^, and carries the CDRs at its tip, which are directly responsible for antigen recognition^16^. The Fc region, by contrast, is composed of two heavy chains also linked by disulphide bonds and mediates interactions with Fc receptors expressed at the surface of immune cells^19,20^ or with soluble cellular effectors^21–23^. These three domains, all built from immunoglobulin folds rich in β-sheets, are joined by a disordered segment known as the hinge, made of two disulphide-linked heavy-chain fragments^11,17^ (Fig. 1). The combination of a flexible hinge, three semi-independent domains, and an overall elongated shape makes full-length IgGs notoriously difficult to resolve by X-ray crystallography^17^. As a result, the individual Fc and Fab domains, smaller and less flexible, have been characterised in far greater detail^23^, and thousands of their structures have been deposited in the Protein Data Bank^24,25^ (PDB). Full-length IgG structures encompassing both the Fabs and the Fc, in contrast, remain rare and are typically resolved at modest resolution (e.g. 5DK3 at 2.28 Å, 1IGT at 2.80 Å, 1MCO at 3.20 Å).

**Figure 1:**
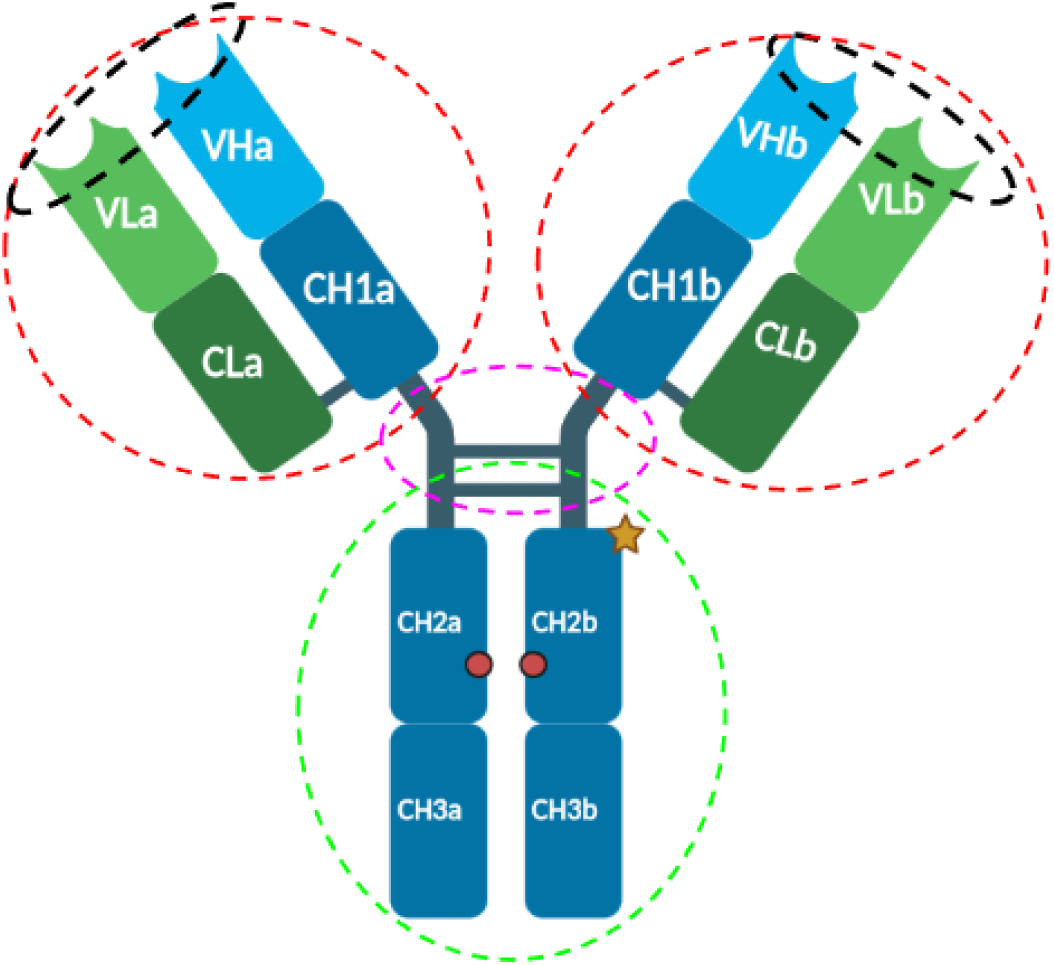
Schematic representation of a full length IgG. The blue regions are the heavy chains. The green regions are the light chains. The green dotted circle represents the Fc region. The pink dotted circle surrounds the hinge region. The red dotted circles are the Fabs, and the black dotted circles are the CDR. The red dots represent the N-glycosylations positions on the Fc. The orange star represents the exposed N-glycosylation position on PMB.

Like most proteins produced by eukaryotic cells, mAbs undergo post-translational modifications (PTMs), and the most prominent of these is N-glycosylation, the covalent attachment of an oligosaccharide to an asparagine (N/Asn/ASN) residue^26^. This attachment occurs at a well-defined consensus sequence, asparagine–X–serine/threonine (where X is any amino acid except proline)^27^. The functional consequences of N-glycosylation are wide-ranging^27–29^: glycan chains can modulate protein folding, thermal stability, function, expression, and pH resistance. In the specific case of antibodies, most attention has focused on the N-glycans localised on the Fc region^15,17,26^, because the Fab domain generally lacks germline-encoded N-glycosylation sites, and only around 15 to 25% of serum IgGs carry one or more N-glycans within their variable regions^30^. Several recent reviews have underlined the impact of Fc glycans on antibody properties^30–35^, and, in particular, on the ability of mAbs to engage the various Fc receptors, an effect attributed to a glycan-induced reduction of Fc flexibility^14,33,36^. This rigidification is very likely what has made most existing Fc structures and the handful of available full-length mAb structures tractable for crystallography in the first place^18,20,22,26^. Building on these observations, glyco-engineering has become an important complementary strategy in antibody design^21,37–40^. Sha et al., for example, showed that controlling fucosylation can substantially enhance antibody-mediated cytotoxicity in the context of cancerous-cell destruction^37^.

A growing body of experimental work has explored how N-glycosylation influences aggregation propensity^34,36^, antigen and receptor binding, and overall antibody stability^34,39–41^. Yet, despite the importance of glycans for antibody engineering, only a handful of *in silico* studies have addressed their effect on the full-length IgG. Molecular dynamics simulations are particularly well suited to fill this gap, since they can provide the atomistic detail needed to rationalise experimental observations across IgG subclasses^14,17,27,32,42^. Among the few computational studies available, most have been limited to isolated Fc fragments^32,36,41,43^, and only a small subset has considered the full-length antibody^33^. Even in this subset, attention has been overwhelmingly directed at IgG1^15^, the most abundant IgG in human serum^42,44^. Other IgG scaffolds exhibit substantial architectural differences and carry distinct oligosaccharides; individual antibodies built on these scaffolds may therefore respond to glycosylation differently from the IgG1 systems studied so far, a possibility we probe here with two antibodies that depart from the IgG1 architecture in different ways.

The present work addresses this gap by studying the effect of N-glycosylation on two full-length mAbs that differ markedly in scaffold: Mab231, a murine IgG2a, and pembrolizumab (PMB), a human IgG4 (Fig. 2). Beyond their different isotypes, the two molecules come from different species, which contributes to the moderate sequence identity between them (Table S1) and to their dissimilar three-dimensional architectures. Both carry two N-glycosylations on their Fc domains, but the glycan patterns differ, both in chemical composition and in spatial location (Fig. 1). PMB is a human IgG4 mAb targeting Programmed-Death 1 (PD-1) in cancer therapy: by blocking the engagement of PD-1 by its ligand PD-L1, it relieves the inhibition of T cells and thereby restores the antitumour immune response. It carries two distinct oligosaccharides on its Fc. Mab231 is a murine IgG2a mAb raised against canine lymphoma cells, and it carries two identical oligosaccharide chains on its Fc.

**Figure 2:**
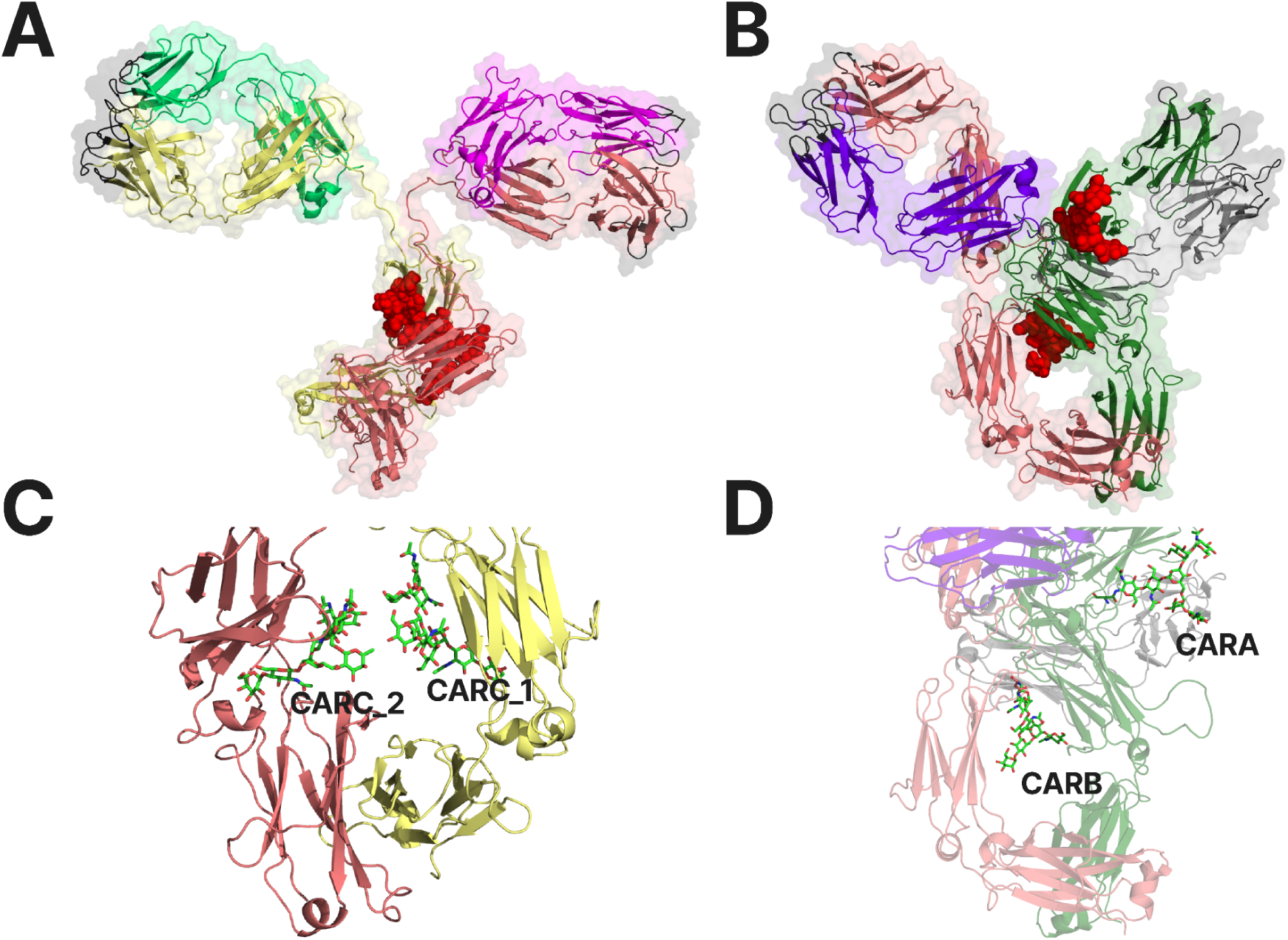
Top: 3D structures of Mab231 (PDB ID: 1IGT) A) and Pembrolizumab (PMB, PDB ID: 5DK3) B), left and right respectively. The chains are colored differently. For Mab231 (A), the pink and yellow chains are the heavy chains. The green and purple chains are the light chains. For PMB (B), the salmon and green chains are the heavy chains. The magenta and grey chains are the light chains. The black zones correspond to the CDRs. In red, are indicated the sugars in space-filling representations. Bottom: Presentation of a “classic” structure of IgG represented by Mab231 (C) and the structural difference of Pembrolizumab (D). The green molecules represent the glycans present in both crystallographic structures.

Both antibodies were investigated by classical molecular dynamics simulations, with and without the full N-glycan chains as described in their deposited PDB structures, in three independent replicates of 1 µs per system, giving in total 12 µs of sampling for all systems. We then performed an in-depth analysis of the impact of N-glycosylation on the accessible conformational space and on the dynamics of the protein, examining a broad set of descriptors, including preferred conformations, flexibility, and local conformational entropy of the residues.

## MATERIALS & METHODS

### Preparation of the initial structures

The crystallographic full-length structures of pembrolizumab (PDB: 5DK3, 2.28 Å resolution, Scapin et al.^26^), a human IgG4, and Mab231 (PDB: 1IGT, 2.80 Å resolution^17^), a murine IgG2a, were retrieved from the Protein Data Bank (https://www.rcsb.org/). The N-glycans considered for our simulations are those resolved in these two deposits, and are represented in Fig. 3 using the *Symbol Nomenclature For Glycans (*SNFG) nomenclature^45^. Their SNFG representation is provided in Fig. 3, and the corresponding CRAM representation in Fig. S1.

**Figure 3:**
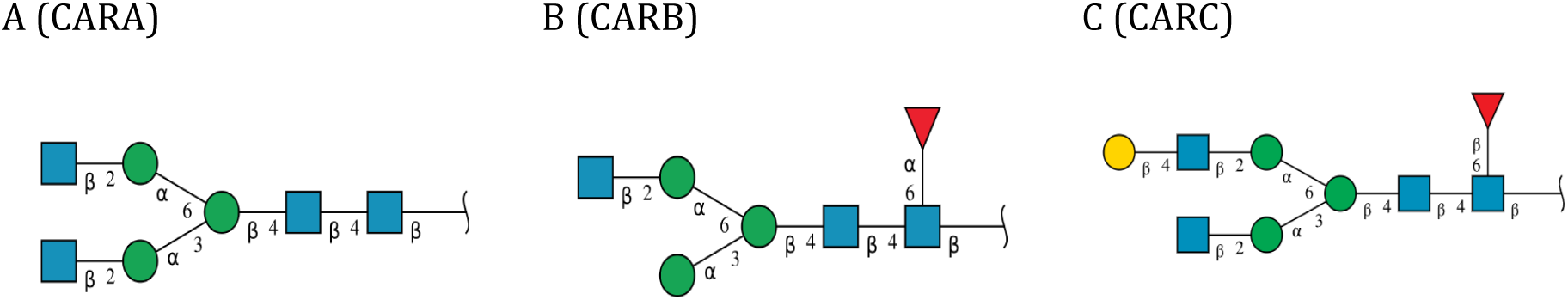
Schematic representation of the carbohydrates. A & B are from the Pembrolizumab, respectively on chain A and B (noted CARA and CARB in the following) Mab231 has twice the carbohydrate C (noted CARC_1 and CARC_2). Blue square stands for N-Acetyl Glucosamine (GlcNAc), green circle for mannose (Man), yellow circle for galactose (Gal), and red triangle for fucose (Fuc). Greek symbols indicate the anomeric form of the hemiacetal/hemiketal. The number at the connection line represents the position of alcoholic hydroxy where the glycosidic bond is formed. This schematic representation is based on SNFG.

The domain delimitations of each antibody are defined in Table 1.

**Table 1:** Delimitations of the fragments for Pembrolizumab and Mab231 structure, with the location of the sugar. Subdomains of FAB-1 and FAB-2 are noted a and b respectively, in addition to the classical labelling. * Chain bound to sugars CARA and CARC_1 to their ASN on PMB and Mab231. ** Chain bound to sugar CARB and CARC_2 to their ASN on PMB and Mab231.

|  |  | 5DK3: PMB (IgG4) |  |  |  | 1IGT: Mab231 (IgG2a) |  |
| --- | --- | --- | --- | --- | --- | --- | --- |
| VL_a | FAB-1 | A | 1-110 | VL_a | FAB-1 | A | 1-106 |
| CL_a |  |  | 109-214 | CL_a |  |  | 114-214 |
| VH_a |  | B | 1-118 | VH_a |  | B | 1-110 |
| CH1_a |  |  | 119-220 | CH1_a |  |  | 120-221 |
| VL_b | FAB-2 | F | 1-110 | VL_b | FAB-2 | C | 1-106 |
| CL_b |  |  | 118-218 | CL_b |  |  | 114-214 |
| VH_b |  | G | 1-118 | VH_b |  | D | 1-110 |
| CH1_b |  |  | 127-220 | CH1_b |  |  | 120-221 |
| CH2_a | FC-1 | B* | 234-341 | CH2_a | FC-1 | B* | 252-357 |
| CH3_a |  |  | 342-446 | CH3_a |  |  | 368-474 |
| CH2_b | FC-2 | G** | 239-336 | CH2_b | FC-2 | D* | 252-357 |
| CH3_b |  |  | 347-444 | CH3_b |  |  | 368-474 |
| CARA | FC-1 | B | N515 | CARC_1 | FC-1 | B | N512 |
| CARB | FC-2 | G | N1177 | CARC_2 | FC-2 | D | N1170 |

### Completion of the structure

The 9 missing residues of PMB, located in the upper hinge region of both heavy chains (P230, A231, P232 on chain B; P230, A231, P232, E233, F234, L235 on chain G), were built with MODELLER^46^ using the automodel option. The default parameters were used. 10 models were considered, and the model with the best DOPE score^46^ was kept. This model was used both for the systems with and without the N-glycans. Mab231 did not need any modelling since it was fully complete.

### Alignment of sequences and structures of both antibodies

The sequences were compared chain by chain (light against light, heavy against heavy) by using the Needleman–Wunsch implementation provided by the NIH website^47^ with default parameters. The structural comparison was carried out with the TM-align^48^ tool, again chain by chain.

### System preparation and molecular dynamics protocol

The CHARMM-GUI^49^ toolbox was used to prepare all four systems (glycosylated and aglycosylated PMB; glycosylated and aglycosylated Mab231). The workflow was identical across systems and proceeded as follows: (i) the protonation state of each titratable residue was assigned at pH 7.4; (ii) the disulphide bonds (16 bonds in PMB and 17 bonds in Mab231) indicated in the SSBND records of each PDB entry were checked and included; (iii) glycosylated systems were built with the Glycan Reader & Modeler tool of the CHARMM-GUI suite, using the crystallographic glycans exactly as deposited (see also the Preparation of the initial structures section); (iv) each system was placed in a cubic solvent box filled with TIP3P water molecules, along with Na⁺ and Cl⁻ ions at a physiological concentration of 0.15 M, plus the additional ions required to neutralise the system; (v) the CHARMM36m force field was used to describe the atomistic interactions for protein and carbohydrates.

The final box dimensions were 16.75 × 16.75 × 16.75 nm³ for glycosylated PMB and 16.76 × 16.76 × 16.76 nm³ for aglycosylated PMB; 17.94 × 17.94 × 17.94 nm³ for glycosylated Mab231 and 17.95 × 17.95 × 17.95 nm³ for aglycosylated Mab231. The resulting systems comprised 481,411 atoms (153,240 water molecules) for aglycosylated PMB and 481,511 atoms (153,039 water molecules) for glycosylated PMB, and 590,198 atoms (189,778 water molecules) for aglycosylated Mab231 and 590,208 atoms (189,640 water molecules) for glycosylated Mab231.

The Charmm-Gui protocol includes a first round of energy minimization consisting of 5000 steps, of the steepest-descent algorithm with a force-threshold convergence of 100 kJ mol⁻¹ nm⁻¹. Bonds involving hydrogen atoms were constrained with the LINCS algorithm, which enabled an integration timestep of 2 fs.

Then, the heating step consists of assigning velocities drawn from a Maxwell–Boltzmann distribution at 300 K. The system was run under NVT and subsequently under NPT conditions, with pressure controlled at 1 bar using the Parrinello–Rahman barostat (isotropic coupling, τ*_p_* = 5 ps, compressibility 4.5 × 10⁻⁵ bar⁻¹) and temperature controlled at 300 K using the Nosé–Hoover thermostat (τ*_T_* = 0.1 ps). The tight thermostat coupling was retained because it was already the CHARMM-GUI default, and no thermal instability was observed at this value. Six successive MD runs were performed as classically proposed by the CHARMM-GUI protocol (the so-called “equilibration” step), each frame being retrieved every 10 ps.

Across these six steps, amounting to 125,000 integration steps at a 2 fs timestep (250 ps in total), the harmonic position restraints on the protein and on the glycan heavy atoms were progressively reduced from 4000 to 0 kJ mol⁻¹ nm⁻². Each fully relaxed system was then run for approximately 10 ns of unrestrained pre-production. The production phase was subsequently carried out without any restraint for 1 µs per replicate using GROMACS 2023.3. For each of the four systems, three independent replicates were generated by drawing different sets of initial velocities from the Maxwell–Boltzmann distribution, bringing the cumulative production sampling to 3 µs per system, so 12 µs in total.

Long-range electrostatics were treated with the Particle-Mesh Ewald method, with a real-space cutoff of 1.2 nm and a grid spacing of 0.12 nm. The van der Waals interactions were treated with a force-switch scheme between 1.0 and 1.2 nm. The neighbor list was updated every 20 steps with a Verlet cutoff of 1.2 nm. Trajectory frames were recorded every 10 ps for all systems, resulting in 100 000 frames per replicate.

All analyses below were performed on the trajectory after the first 200 ns, taken as the onset of the conformational plateau from the RMSD (Fig. S4). The trajectory-wide descriptors (RMSD, RMSF, Neq and contacts) used 80 000 frames per replicate. For the Gaussian-mixture clustering of the Fab-orientation angles, the three replicates of each glycoform were pooled and randomly subsampled (fixed seed) to 9000 frames per glycoform, giving a balanced 18 000 frames per system. Cluster populations (Figs. 6 & S18 & S19) therefore sum to 18 000 for each antibody.

### Protein Blocks

The Neq is defined as a conformational entropy that quantifies the diversity of local backbone conformations sampled by a residue over the trajectory. It is derived from a structural alphabet of 16 letters (*a* to *p*), the Protein Blocks (PBs)^50^, each PB representing a canonical 5-residue local backbone conformation defined by the φ,ψ dihedral angles. Every frame is translated into a string of PB letters, with each letter assigned to the central residue of its 5-residue window. The four terminal residues at each end of a chain are assigned the letter *Z* for completeness (with no physical meaning). For each amino acid, the frequency fₓ of every PB letter *x* is computed over the whole trajectory, and the Neq is then calculated as a Shannon-entropy-based index (Equation 1). The Neq ranges from 1 to 16, for when the residue adopts a single state throughout the simulation and when the residue explores the entire structural alphabet, respectively. Values of Neq can be identified as 4 different classes: a rigid position at Neq = 1, flexible at Neq > 4, highly flexible at Neq > 6, and extremely flexible just like a disordered region at Neq > 8. This analysis was realized for each simulation not pooled, it was realized per replicas.

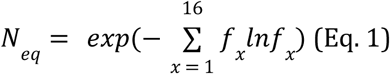

### Geometrical Analyses of the conformations

Antibodies are frequently described through a small set of characteristic shapes^15^, the T-shape, Y-shape, and the less common λ-shape, defined primarily by the relative orientation of the two Fabs regarding the Fc (Fig. S2). To capture this orientation quantitatively, we used the θ and φ angular descriptors originally proposed by Saporiti et al^15^. The origin of the coordinate system was placed at the centre of mass of the hinge region (CoM-Hinge). Two alternative definitions of the axes were tested: (i) the z-axis defined as the vector from the CoM-Hinge to the joint centre of mass of the two CH2 domains, and the x-axis defined as the vector from the CoM-Hinge to the centre of mass of Fab1 (or Fab2); (ii) the z-axis and x-axis defined instead as the principal inertia axes of the Fc and of the Fab, respectively. In both cases, the y-axis was taken perpendicular to the (z, x) plane. The angles θFab-1 and θFab-2 were then defined as the angles between the z-axis and Fab-1 or Fab-2, and the θ inter-Fab angle as the angle between the two Fab inertia axes (Fig. S2.A). The molecular dynamics simulations were centered on the Fc of the mAbs and considered as a reference for measurements.

We also examined the elbow angle, which captures the potential crease between the variable and constant regions of the Fab and provides a measure of intra-Fab rigidity. Following Stanfield et al.^51^, it was defined as the angle between the VH → VL and CH1 → CL centre-of-mass vectors (Fig. S2.B). Finally, we measured the dihedral hinge angle describing the twist of the two Fab arms around the long Fc axis. All definitions are provided in the supplementary material (Fig. S2.B & S2).

### Clustering of the Fab orientations

The two-dimensional (θ, φ) orientation of each Fab arm was clustered with a Gaussian mixture model^52,53^ (GMM) using the method *full* for the covariance. Prior to clustering, each (θ, φ) pair was mapped onto the unit sphere as a three-dimensional cartesian vector so that the angular periodicity of φ and the pole behaviour of θ are handled consistently. Mixtures with full covariance matrices were fitted for a number of components k from 1 to 10, each with five independent initialisations, and the optimal k was selected as the one minimising the Bayesian information criterion (BIC). Each frame was then assigned to its most probable component, and the medoid of each cluster, defined as the frame closest to the cluster mean, was retained as its representative conformation. All the angles were clustered independently.

### Statistical Analysis

Each descriptor was compared between the glycosylated and aglycosylated states replicate by replicate, and the effect sizes and overlap coefficients reported in the Supp. Mat. Results (Table S3 & S4) are the medians of the three per-replicate values. We did not concatenated the replicates for this comparison, because they sample different regions of the Fab–Fc space: pooling would then blend an inter-replicate difference into the glycosylated-versus-aglycosylated comparison and could make the two states look more distinct than they are within any single replicate. The angular descriptors θFab-1, θFab-2 and θInter-Fab between the two forms for each antibody were compared with the Mann–Whitney U test. These angle values were also used to assess convergence, via the Flyvbjerg–Petersen block-averaging procedure (Fig. S3 and associated comments).

### Analysis of contacts between amino acids and carbohydrates

Contacts between residues, between residues and the glycans, were computed with the MDAnalysis^54,55^ Pyresidues and. A contact between two amino acids was defined when the distance between their Cα atoms was less than or equal to 8 Å. The same 8 Å criterion was used for residue–glycan contacts, but computed between the Cα of the residue and any heavy atom of the glycan chain. For each contact, we report its persistence, defined as the percentage of analysed frames in which the contact is present, pooled across the three replicates of each system:

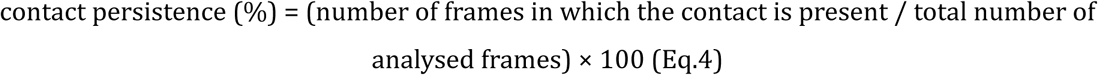

Contact percentages reported in the Supplementary Material and briefly reported in the main text correspond to the fraction of frames in which a given contact is observed, computed per frame after equilibration and pooled across the three replicates of each system.

### Allosteric network

The effect of glycans on allosteric communication was examined with the allopath tool developed by Delemotte and co-workers^56,57^ (https://github.com/delemottelab/allosteric-pathways), with the glycans as the source of the network and the antibody residues as the sink. Mutual information was estimated with the bootstrap set to 10 and n_blocks_col, n_blocks and n_chains each set to 1. The first 200 ns of each replicate were discarded, and the three replicates of each glycosylated antibody were merged for the analysis.

## RESULTS

### The mechanical nature of the antibodies induces large conformational changes

Along the dynamics, the two proteins strongly deviate from their initial conformation, with a global calculated RMSD for the full-length antibody reaching 20-25 Å (Fig. S4). This corresponds to large conformational changes, slightly smaller for the glycosylated forms. Actually, the large values result from changes in the relative positioning of the Fc and Fab domains, which are not significantly deformed by themselves (Fig. S4).

The analysis of contacts along the simulations shows that some native contacts are lost but compensated by the formation of new contacts for both antibodies (Fig. S5). The presence of N-glycosylation tends to slightly limit the loss of native contacts for PMB (95% with sugar versus 88% without sugar). In the case of Mab231, the number of native contacts remains very similar regardless of the presence or absence of sugar (∼95%). Overall, contacts within subdomains are mainly maintained, while new contacts are formed between the subdomains, induced by their relative motions. This shows that both proteins remain stable, maintaining their secondary structures and preserving most of the native contacts despite important conformational changes.

In summary, these changes reflect the mechanical character of antibodies, namely rigid subdomains that move freely, thanks to the highly flexible hinges that connect them (see RMSF analysis, Fig. S6). To better understand the local flexibility, especially these highly flexible hinges, we considered an original description that follows the backbone dihedrals distribution, namely the Protein Blocks (PB). This description allows both the characterization of the protein backbone conformations through a finite number of discrete states and a quantification of their diversity along an MD trajectory through the Neq measure. The Neq is defined as the equivalent number of PBs, for every residue of both antibodies in both glycosylation states. Importantly, the Neq is insensitive to any choice of the reference for the superimposition.

For both antibodies and whatever the glycosylated state, the distribution of PBs confirms the stability of the ⍰-sheet domains that are described with the PBs *d* and *e*. For Mab231 (Fig. 4) in these regions, the residues transit mainly between these two states (Figs. 4.A & 4.B). Interestingly, the frequency associated with a given PB rarely equals a high value. This means that, along the simulation, the residues explore different states, as confirmed with the Neq values (Fig S7 & S8). This is particularly true for some residues of the hinge regions where the Neq can reach average values equal to 4 and even 6 for some residues (Figs. 4.C & 4.D), which demonstrates a greater sampling of the conformational for the hinge region. Outside the hinge domain of Mab231, (Figs.4A, S9 & S.10) despite the presence of the glycans in the Fc region, we do not observe any major differences either between the number of visited states or the types of states, compared to the same region without the glycan. In contrast, for PMB (Fig. 5), the residues of the - sheets are mostly represented by PB *d* and transit extremely rarely in other PBs, even to the closest one (Figs. 5.A & 5.B). Conversely, more residues outside these regions are adopting various states (Figs. 5.A, S9 & S10), e.g. residues 420 (CH1a domain), 430 (CH1a domain), 490 (CH2a domain) and residues 1082 (CH1b domain), 1092 (CH1b domain), 1152 (CH2b domain). The Neq can reach 8 (Fig. 5.C & 5.D), specifically in hinge domain H2, indicating a higher flexibility compared to the equivalent regions in Mab231. This increased flexibility of hinge H2 might be related to differences in the orientation and type of the sugars close to hinges H1 and H2.

**Figure 4:**
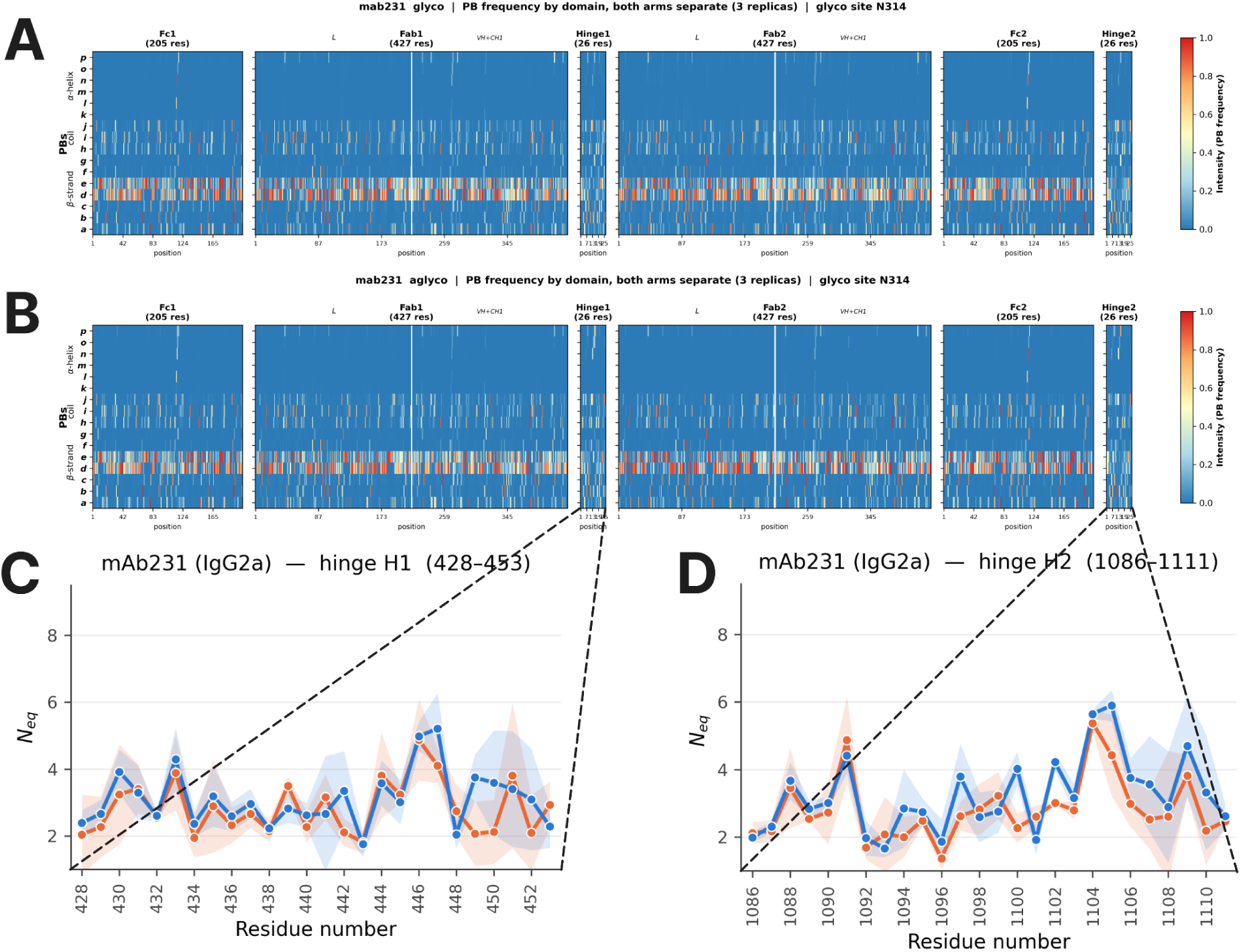
Map PB for glycosylated (A) and aglycosylated (B) Mab231, the intensity of presence of the blocks is represented by the bar color at the right of the figures, and Neq (C and D) values associated to the hinge domains, the aglycosylated simulations are in orange the glycosylated ones are in blue, the blur representation is the standard-deviation.

**Figure 5:**
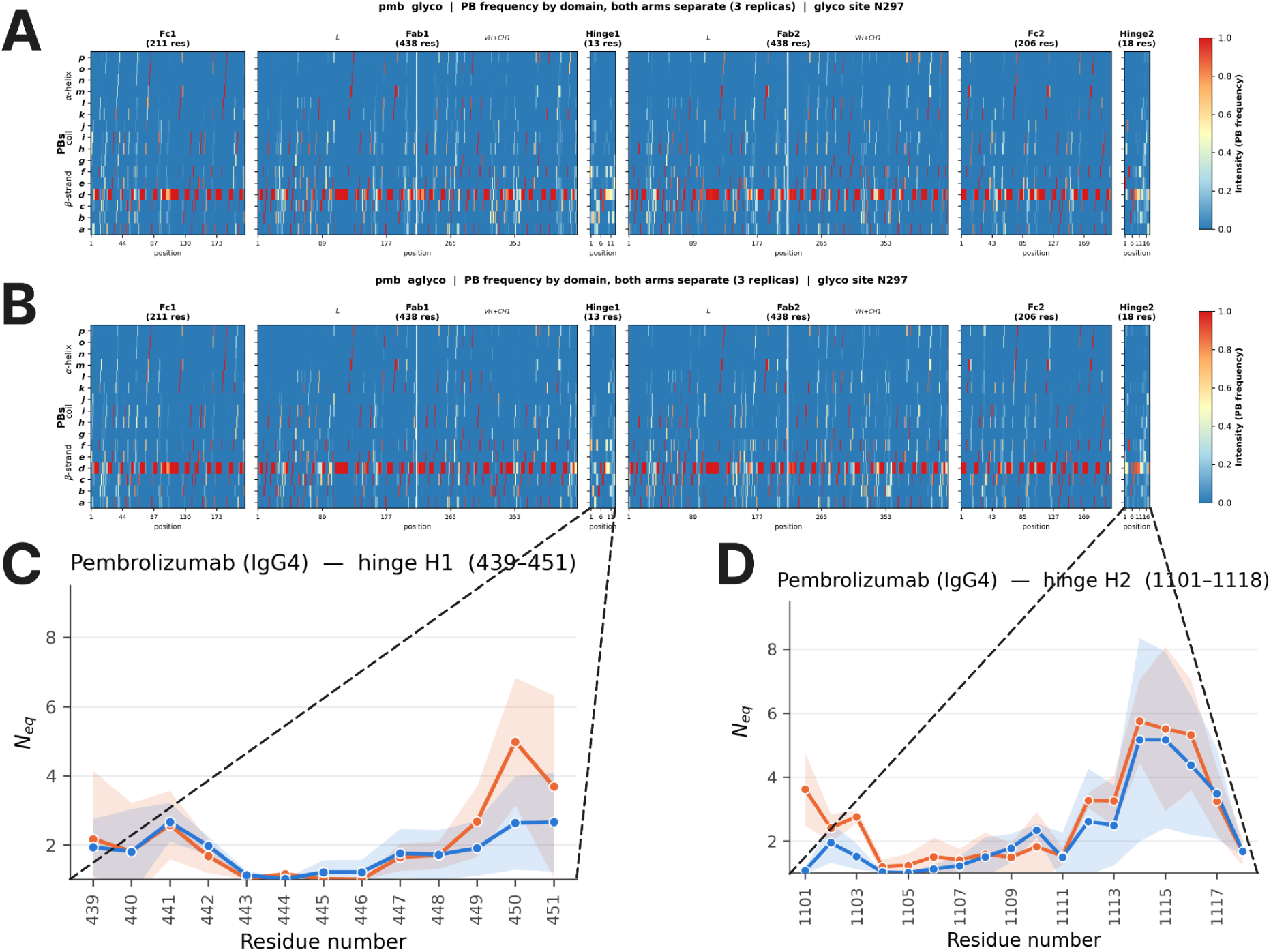
Map PB for glycosylated (A) and aglycosylated (B) PMB, the intensity of presence of the blocks is represented by the bar color at the right of the figures, and Neq (C and D) values associated to the hinge domains, the aglycosylated simulations are in orange the glycosylated ones are in blue, the blur representation is the standard-deviation.

**Figure 6:**
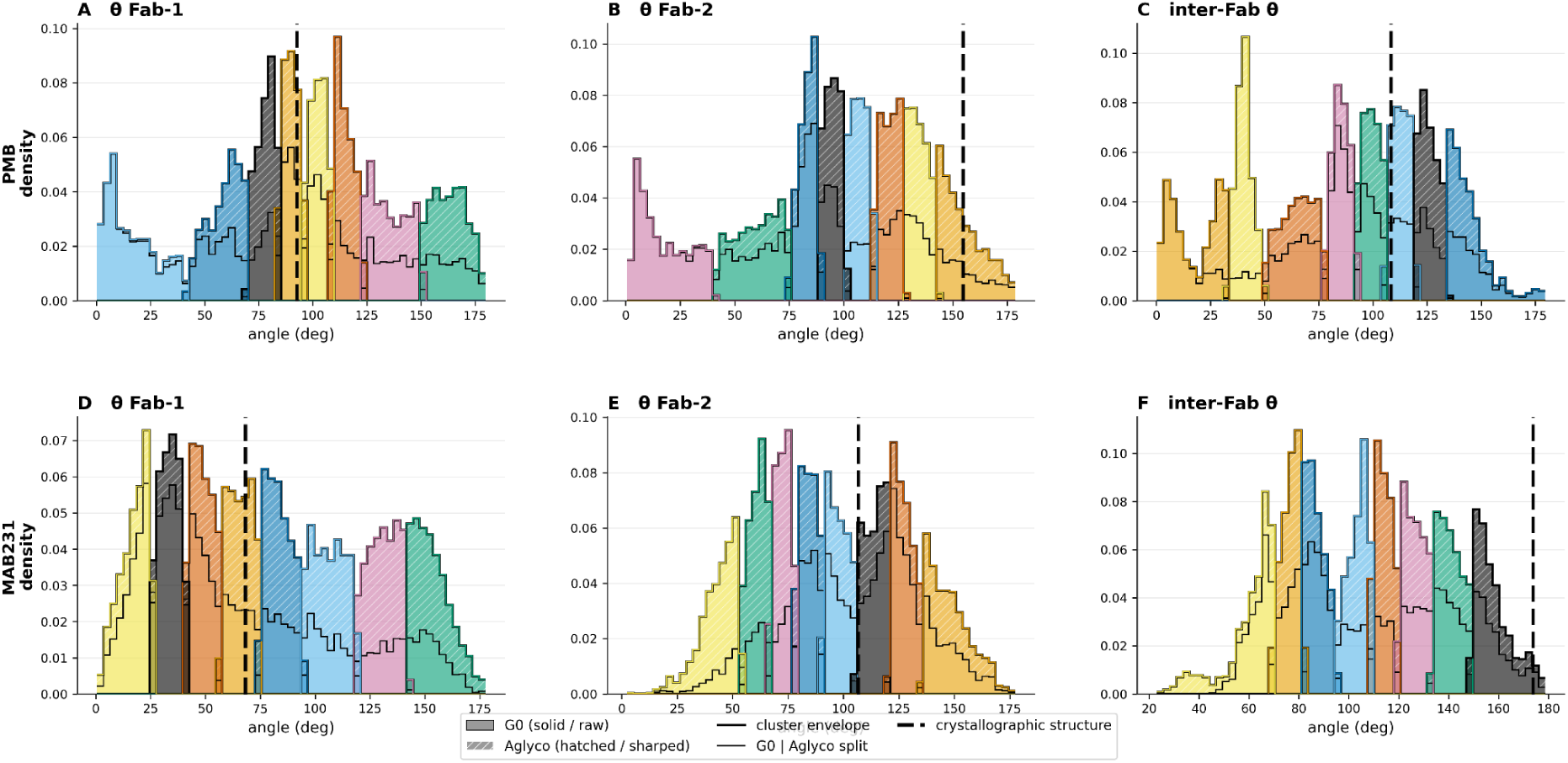
GMM clustering of the angle values adopted by the Fabs for PMB (A, B, C) and Mab231 (D, E, F). The aglycosylated (Aglyco) population is represented with dashes while the glycosylated (G0) is full. Both are separated by dashed black lines on the distributions. Each cluster is represented by different colors. The crystallographic structure is represented by a dotted black line in the distributions.

Although substantially longer simulations would be required to explore the full conformational landscape, the present simulations reveal important features of Ig dynamics over the accessible timescale. In particular, the hinge residues sample distinct torsional states that drive large-scale conformational changes, apparently independent of the glycosylation state. These results further show that even small differences in these mechanically important regions can markedly affect the relative positioning of the rigid domains and, consequently, the contacts they can form.

### Do the sugars impact the conformational space visited by the protein and how?

The analysis detailed above focuses on local properties. But, as we highlight, local changes can have a global impact on the conformations adopted by the protein. Consequently, we considered geometric features that can better define putative global conformational changes.

Among the descriptors previously used to characterise antibody conformations, angles between domains were considered as relevant to better describe potential specific structures (Figs. S18 & S19 & S20). Consequently, we defined a set of angles that capture the positioning of the Fab domains with respect to the Fc domain: θFab-1 and θFab-2 are angles between the Fab-1 and Fab-2 axes and the Fc axis respectively, θinter-Fabs is the angle between the Fab-1 and Fab-2 axes, and φFab-1 and φFab-2, the rotation angles of Fab-1 and Fab-2 around the z axis (Fig. S2). We merged all the conformations from the two states and examined the distributions of the angles. For all angle types, the values span a wide range, confirming broad conformational sampling. The distribution of each angle was analysed using GMM clustering method, (Fig. 6 & S18 & S19). According to the Bayesian Information Criterion *BIC*^52^,the optimal clustering defined by the lowest BIC among the values tested from (k=1) to (k=10), corresponded to (k=8) clusters. In most cases, the clusters are populated, both with glycosylated and aglycosylated conformations.

Despite the slow relaxation indicated by the blocking analysis (Fig. S3) and the global RMSD drift (Fig. S4), GMM clustering of the Fab angular descriptors (Fig. 6) reveals a consistent conformational landscape for both systems. The aglycosylated and glycosylated antibodies broadly sample the same angular range [0°;–180°]. Thus, N-glycosylation does not appear to substantially modify either the nature or the multimodal distribution of the interdomain motions. The main difference is observed at small angles, which are slightly more populated in the glycosylated antibody; this is also represented by the conformations presented in Fig. S20. The clusters compositions are presented in Table S2.

These changes in angles contribute to relative motions between domains that can be also observed thanks to the DCCM (Fig. S11 & S12 & S13). Indeed, for both proteins, the analysis of the DCCM confirmed i) the existence of rigid domains, as shown by the correlated motions (yellow squares), separated by hinge region ii) anticorrelated motions between some subdomains, e.g. CH1a in FC1 vs VLb in Fab1 or CH2a in Fc-1 and VHb in Fab-2 for PMB, or VHa in Fab-1 with CH2a in Fc1 and VHb in Fab-2 and CH2b in Fc-2 for Mab231 (Fig S11). These motions involving Fab-1 versus Fc-1, or Fab2 versus Fc-2 or Fab-1 versus Fab-2, explain the large range of values covered by the angles detailed above. The values are in general smaller for Mab231 compared to PMB. The absolute difference between the glycosylated and aglycosylated matrices (Figs. S11 & S12) show changes between many differing pairs, particularly for PMB, the largest apparent changes being between the Fc and the Fabs, for example CLb relative to CH2a and CH2a relative to CH1b. However, considering the variation between replicates (Fig. S13), these differences cannot be fully attributed to the glycosylated state.

### Do and how do the sugars interact with the protein residues ?

The analyses presented above indicate that the carbohydrates do not significantly alter the overall dynamics of the two antibodies, but that they can modulate the conformational regions explored, and in particular the relative orientation of the Fab and Fc domains. Such modifications could in turn influence the ability of the protein to interact with its partners, and thereby affect its functional properties (Figs. S20 & S21).

To examine the potential impact of the sugars on the partner interacting regions, the so-called shielding effect, we identified the residues in contact with each carbohydrate and their persistence during the simulation (Figs. 7 & 8 & S22 & S23). The regions involved in these sugar protein interactions are mapped onto the structures in Fig. 8 for easier visualisation.

**Figure 7:**
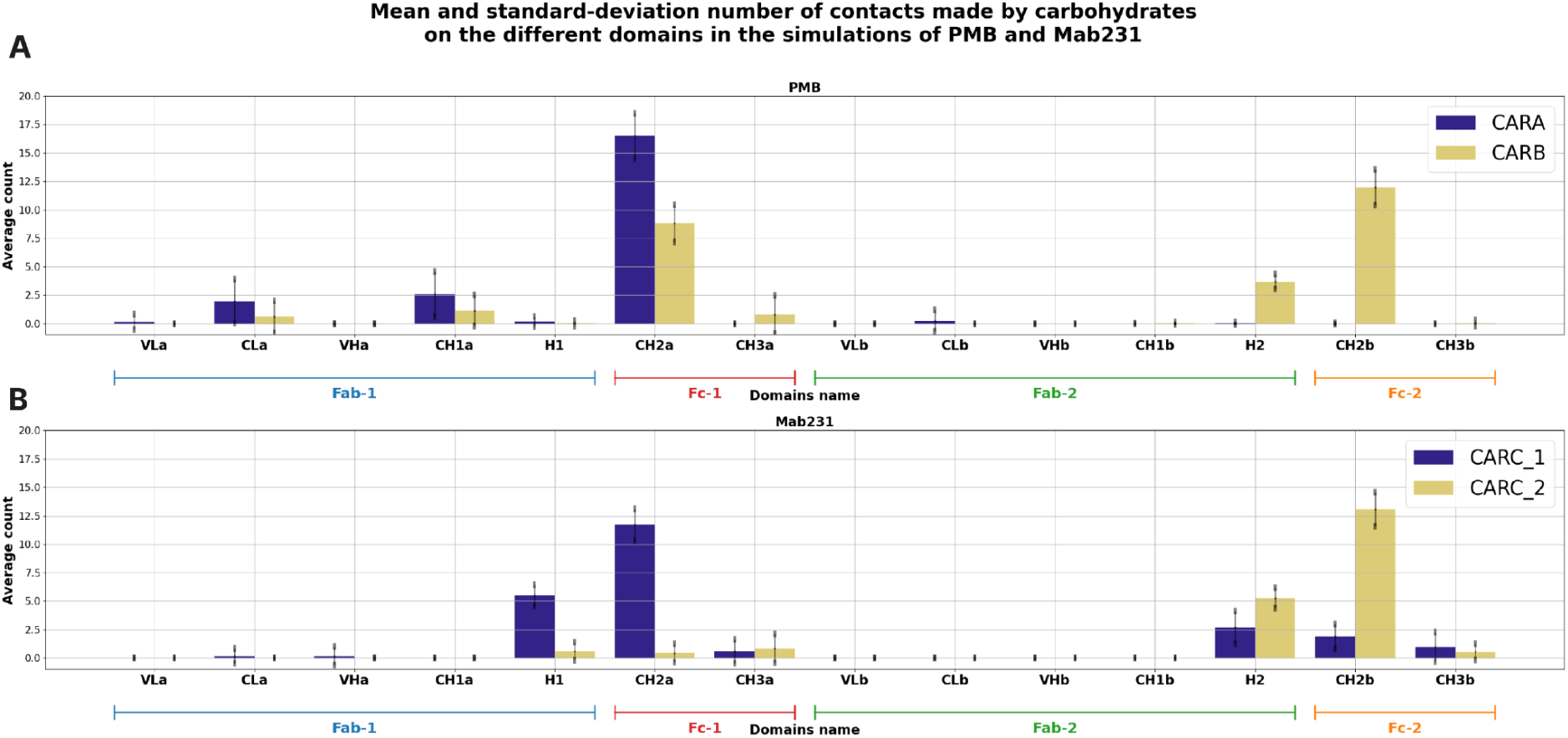
Average number of residues in interactions with the different carbohydrates in the simulations of PMB (A) and Mab231 (B). Both antibodies have two glycans, CARA from PMB in blue and CARB from PMB in orange. CARC_1 from Mab231 in blue and CARC_2 from Mab231 in orange. The black lines on each bar are the standard-deviation values.

**Figure 8:**
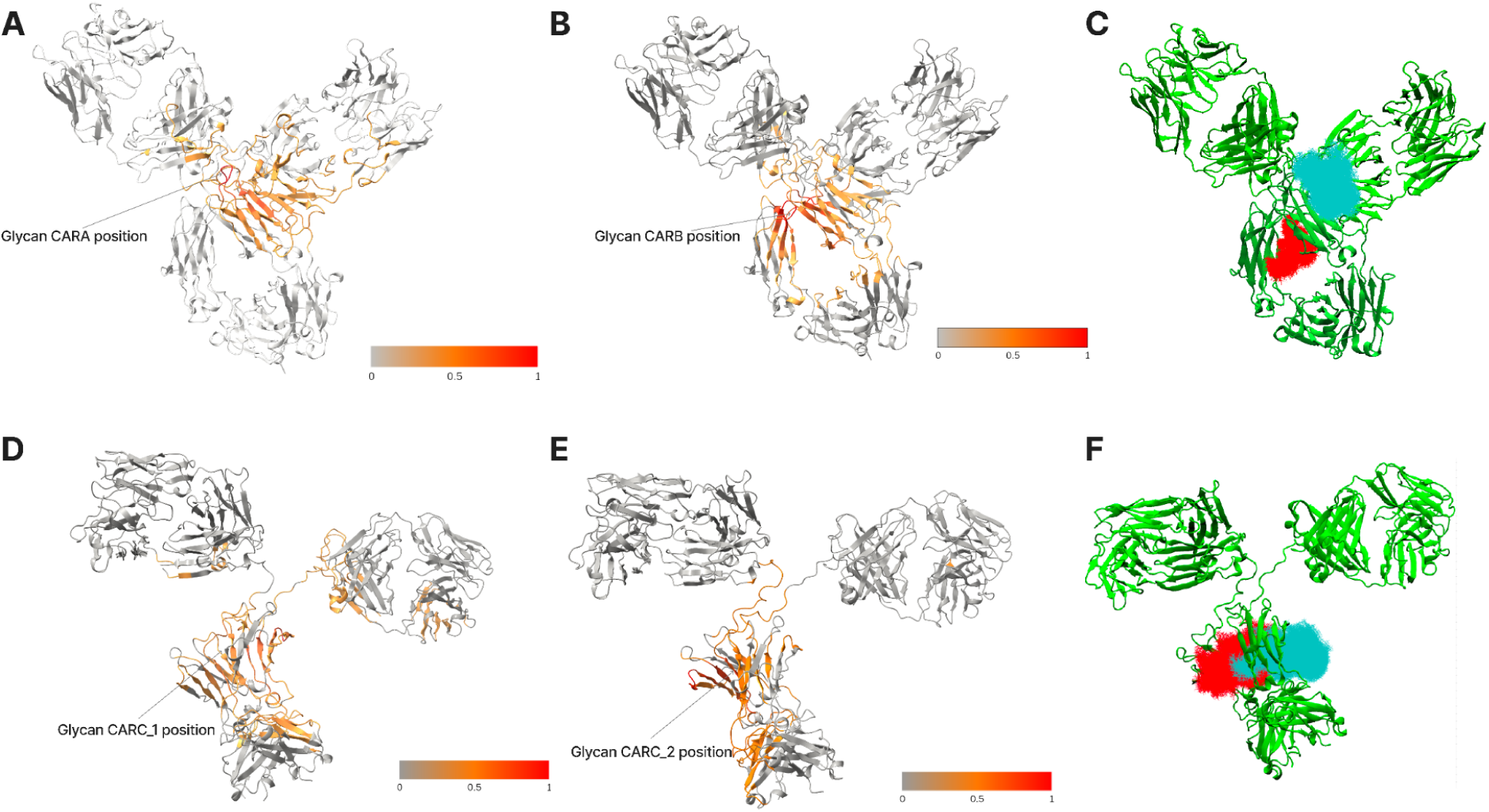
The interactions observed during all the simulations are projected on the crystallographic conformation for each mAb. Top) PMB residues in contact with A) CARA sugar and B) CARB sugar C) ensemble of conformations of the sugars CARA in cyan and CARB in red. Bottom) Mab231 residues in contact with D) CARC_1 sugar and E) CARC_2 sugar; F) ensemble of conformations of the sugars for CARC_1 and CARC_2, cyan and red respectively. The color bar represents the frequency of sugar contacts with the considered residue, from 0 (never in contact, in grey) to 1 (the most involved in contact with the glycan, in red).

For PMB, the highest number of contacts occurs between CARA and the CH2 residues of Fc-1, and between CARB and the CH2[a/b] residues of both the Fc-2 and the Fc-1 domains (Fig. 7.A). Each glycan therefore interacts primarily with the subdomain to which it is covalently bound (Fig. 7). Interestingly, a small number of contacts also occur between CARA and certain Fab residues (Fig. 7.A). This is also observed for CARB to a lesser extent (Fig. 7.A). This is explained by the large motions of PMB during the simulation (Figs. S20 & S21), which bring Fc-1 and Fab-1 into closer proximity, combined with the larger conformational landscape accessible to CARA because of its specific location (Fig. 8). CARB, in contrast, is sequestered between the two Fc fragments (Fig. 2) and can only rarely reach the Fab regions. Beyond their spatial location, the chemical composition of the two glycans (Fig. 3) may also contribute to these differences, as reported in previous studies^31,58^. Finally, a few contacts occur between the two sugars and the hinge regions.

Mab231 behaves in a comparable way (Fig. 7.B): each carbohydrate interacts mostly with its own domain and with the facing Fc fragment. The two sugars show a similar set of interactions, as expected from their identical chemical composition and symmetric locations. Both can also reach the Fab regions to a lesser extent than carbohydrates of PMB.

Persistence of most contacts is quite high, even with Fabs residues (Figs. S22 & S23). Post-translational modifications, and N-glycans in particular, have actually been reported to act distantly from the modification site^59,60^. Long-range communication between the Fc and the Fab has been described in an aglycosylated IgG4^61^. To probe this, we computed the allosteric network with AlloPath^56,57^, taking each glycan in turn as the source (Fig. 9).

**Figure 9:**
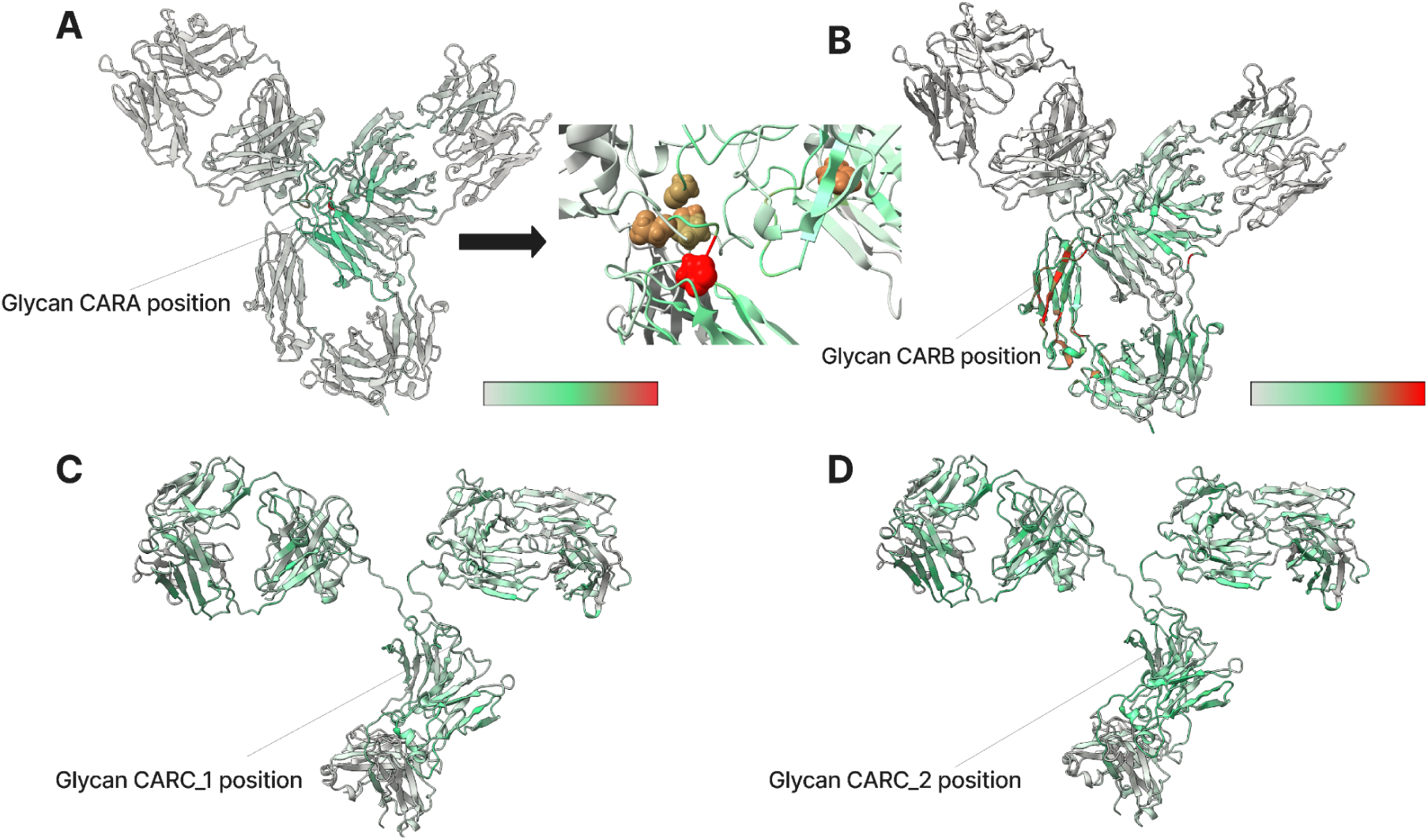
Projection of the implications of the residues in the allosteric network with the glycans as source. A) CARA on PMB. B) CARB on PMB. C) CARC_1 on Mab231. D) CARC_2 on Mab231. The color goes from grey to cyan to red. It illustrates how involved is the residue in the allosteric network of the mAb.

For PMB, the network sourced on CARA (Fig. 9.A) extends well beyond the Fc, reaching the hinge and the Fab, where several residues are strongly involved. This correlates with the contact analysis, which found direct CARA–Fab interactions enabled by the conformational freedom of this glycan. For CARB (Fig. 9.B), the network is concentrated on the Fc fragment carrying the glycan, with a smaller Fab contribution like in the contact analysis.

For Mab231, the two glycans (Fig. 9.C & 9.D) behave similarly, their networks also reaching the variable regions of the Fabs. The individual contributions are weaker than in PMB but cover a larger part of the structure. Consequently, the glycan-sourced networks extend beyond the Fc and into the Fabs in both antibodies. The CDRs are not involved, but the core of the Fab domain participates in the case of Mab231: these are framework residues, known to influence the conformation of the hypervariable loops and hence antigen recognition^62,63^. The networks also reach the hinge, consistent with the differences in overall shape between the two systems, and such a reorganisation of the domain arrangement could in turn affect how the antibodies engage their antigens and receptors.

## DISCUSSION

To date, most theoretical and experimental studies have focused on the most abundant IgG class, IgG1, and have generally restricted their exploration to the Fc domain, which carries the glycosylation sites and mediates receptor binding. As a result, the influence of Fc glycosylation on the Fab domains has rarely been addressed. To our knowledge, it is even rarer for antibodies built on non-IgG1 scaffolds such as the two studied here, which differ appreciably from IgG1, in particular in the hinge region connecting the Fab and Fc subdomains.

In the following sections we therefore discuss the few available studies on the two antibodies examined here in light of our own results, before turning to the more abundant literature on IgG1.

The first molecular dynamics simulations of the Mab231 murine IgG2a were carried out by Wang et al^59^, who examined a range of systems, from the full-length glycosylated protein down to individual subdomains, over trajectories limited to 40 ns. For the full-length systems, they reported similar backbone fluctuations in the glycosylated and aglycosylated states, except within the CH2 domains. When the CH2 domains were isolated from the rest of the mAb (CH3, Fabs and hinge), however, the loss of glycans markedly reduced their structural stability, reminiscent of thermal stress. The secondary structure was preserved, but the tertiary and quaternary organisation was perturbed: the two CH2 domains shifted relative to one another in the absence of glycans, exposing to solvent a region the authors described as aggregation-prone. They interpreted this as the sampling of a more “open” form of Mab231, thought to be required for Fc-receptor binding. More recently, Ma^64^ investigated glycan dynamics in Mab231, together with the potential influence of antigen and receptor binding. This work focused mainly on distances between the glycans themselves and between the carbohydrates and the CH3 domains, and found that the presence of binding partners altered these distances only slightly.

In our study, which extends over a longer timescale, we observed an equilibrium between two distinct, well-separated distance distributions (Fig. S24), which suggests that the intrinsic dynamics of the glycosylated antibody alone may be sufficient for it to access the “open” form invoked in several studies, should that form indeed be required for receptor binding. This interpretation, however, hinges on how the distance used to characterise the different Fc conformations is defined, a choice that is far from straightforward. In IgG1, residue P238 is commonly used to define the aperture, but this residue is absent in this murine IgG2a, and it also lies within the flexible hinge, which further complicates its use as a reference.

For the IgG4 Pembrolizumab (PMB), Natesan et Agrawal^61^ compared its dynamics with that of an IgG1 using implicit-solvent molecular dynamics, assigning both systems the same A2G0 glycan, the most simplified glycan model^58^. Their study did not address glycosylation as such, focusing instead on how the starting conformation shapes the subsequent dynamics. The main point of overlap with our work is that their IgG4 model, like our PMB, sampled a wide diversity of Fab–Fc arrangements.

Spiteri et al.^58^ combined several experimental techniques with Monte Carlo (MC) simulations to probe the role of Fc glycosylation in one human IgG4, the A33. In solution X-ray and neutron scattering, they found only minor differences between the glycosylated and aglycosylated states in the radius of gyration and in the relative distances between the Fab, Fc and Fab-Fc subunits, indicating that the two states are barely distinguishable at the macroscopic level. The neutron data, however, pointed to greater conformational freedom in the aglycosylated form. Building atomistic models with MC sampling, they generated large ensembles consistent with the experimental data and confirmed that the Fc domain accesses a broader range of conformations when aglycosylated. This points in the same direction as our Protein blocks analyses and the putative impact of glycans interactions with the H1 hinge region, which decreases its local flexibility. It is also correlated with our angular analysis (Fig. 6), in which the orientation of the Fabs relative to the Fc tends to get closer or the get more distant, between glycosylation states.

Although they did not consider glycosylation, Tarenzi et al.^65^ studied aglycosylated pembrolizumab both without (apo) and with (holo) its target PD-1, using all-atom molecular dynamics, so their apo-state results provide a natural point of comparison with our aglycosylated PMB. Clustering their trajectories by structural RMSD, they obtained five clusters, each defined by a distinct set of contacts among Fab1, Fab2 and Fc. Across all clusters, they identified interaction networks linking residues in the Fc to residues in Fab1 or Fab2 along a pathway running through the hinge. A comparable network emerges from our own data suggesting it may be a recurring feature of PMB across these independent studies.

### Comparison with an IgG1 study

As noted above, most studies have targeted IgG1, and only a handful have considered the full-length antibody. Zhang et al.^36^ examined how glycosylation shapes the conformational dynamics of an IgG1 Fc domain, showing that removing the glycans on the CH2 domains allow the two chains to get closer over the course of the simulation, whereas the glycans maintain a larger inter-CH2 separation. For Mab231, our fluctuation data cannot tell whether the intrinsic flexibility of the CH2 domains changes with glycosylation. The distance between the two CH2 domains (calculated on the center of mass), however, did vary with the glycosylation state (Fig. S24.B). The glycosylated form populates two states, both slightly more compact than the X-ray structure: a more open one (∼37 Å) and a more closed one (∼33 Å). The aglycosylated form, by contrast, populates a single state, at distances intermediate between the two. Because our longer trajectories sampled both the closer and the more open arrangements in the glycosylated system, this behaviour may reflect a “breathing motion” of the Fc domain in Mab231.

In a related vein, Subedi and Barb^66^ used NMR spectroscopy on an isolated IgG1 Fc fragment to probe the effect of N-glycosylation. They found that it favours the Fc-receptor interaction: rather than altering the Fc quaternary structure, glycosylation stabilises the C′E loop that carries the glycans (residues 280 to 297 in IgG1, corresponding to the equivalent region in PMB) and forms part of the binding interface with the receptor FcγRIIIa. They reported a strong correlation between the conformation of this loop and receptor affinity. Our full-length simulations of PMB and Mab231 likewise show no major effect of the glycans on the quaternary structure (Fig. S20). Our flexibility metrics, dominated as they are by replicate-level offsets, cannot resolve changes localised to a single loop such as C′E, so we can neither confirm nor contradict their observation. Beyond the differences in Fc sequence, the presence of the Fabs and of the hinge in our systems may also contribute to this divergence between the two studies.

Among the very few studies addressing glycan effects on a full length antibody, Saporiti et al.^15^ offered important insights into the role of the sugars in adalimumab, an IgG1. They built the complete system as a homology model combining two X-ray structures, and examined three variants: glycosylated afucosylated, glycosylated fucosylated and aglycosylated. As in our work, they ran long molecular dynamics simulations of 1 µs with three replicates per model in explicit water. They found the CH2 flexibility to be broadly similar regardless of the presence of glycans, except at the very beginning of the region, which is disordered and highly flexible because it connects to the hinge, and which stood out in the glycosylated afucosylated form. Turning to the opening and closing of the Fc, they reported that the absence of sugar, or the presence of fucose, was associated with a more compact, apparently closed Fc, using the distance between the two glycan bearing asparagines as the probe. In their interpretation, these structural differences relate to the capacity of adalimumab to engage its receptor, in particular around the CH2 region, the afucosylated glycan tending to preserve an open Fc and to leave residues more solvent exposed and available for interaction, in line with the work of Wang et al^59^.

Our own data do not reproduce this picture in a straightforward way, and the analysis is descriptor-dependent. Indeed, using the asparagine bearing glycans for measuring CH2 distance, the glycosylated form of PMB samples a narrow, unimodal distribution centred near 33 Å, whereas the aglycosylated form is markedly broader and bimodal, populating both a more compact arrangement near 28 Å and a more open one near 39 Å (Fig. S24.A). The glycans therefore restrict the range of Fc arrangements explored rather than holding the domains apart. For Mab231, which carries a fucose on both glycans, the two distributions are essentially superimposable (Fig. S24.B), so no comparable effect can be discerned.

When the distance between the centers of mass of the two CH2 domains is used instead (Fig. S24.C and S24.D), the picture changes again, and in a way that is consistent across the two antibodies: the glycosylated form is centred at a smaller CoM distance than the aglycosylated one, by roughly 1 Å in both PMB (35.2 Å against 36.4 Å) and Mab231 (34.7 Å against 35.4 Å), and it is again the narrower of the two distributions. Relative to the crystallographic structure, the glycosylated PMB sits essentially at the X-ray value, whereas for Mab231 both simulated states are more compact than the deposited structure.

Taken together, these observations show that the influence of the glycans on Fc opening cannot be settled unambiguously in either antibody, because the answer depends on the reference chosen. On the asparagine criterion the glycans mainly narrow the range of arrangements sampled, while on the CoM criterion they are associated with a slightly more compact Fc, which is the opposite of what would be expected if Fc glycans systematically promoted CH2 opening. This is a reminder of how difficult it is to characterise open and closed states when the structural references differ markedly between antibodies.

In summary, the angle distributions of PMB and Mab231 differ slightly between the two glycosylation states, but not reproducibly across replicates, so we do not interpret them as glycan effects. What does hold is that the glycans reach beyond the Fc to contact Fab residues, and that the glycan-sourced allosteric networks point to the same Fab framework regions; these contacts could in turn modulate antigen or receptor engagement, a hypothesis that remains to be tested. Subedi and Barb^66^, dissecting the allosteric role of Fc N-glycosylation in an IgG1, likewise observed structural perturbations in loop regions; because their study was confined to the Fc, it could not detect the Fab-associated effects we describe here, which may explain the contrast with our findings.

Finally, it is also important to underline that PMB is a chemically modified antibody, bearing two glycans with a different chemical composition. The crystal 3D structure has trapped an unusual λ shape. However, the present set of simulations shows that the structure in solution departs from it and adopts several conformations, including T-shape and Y-shape, giving some robustness to the conformational landscape visited. Mab231, which bears two identical glycans, also leaves its initial T-shape for visiting different conformations.

## CONCLUSION

In this work we compared how N-glycosylation relates to the behaviour of two full-length monoclonal antibodies, pembrolizumab, a human IgG4, and Mab231, a murine IgG2a, using all-atom molecular dynamics simulations. The two molecules differ in their glycan patterns, their sequences and their conformational features, which let us ask which features of the subdomain cross-talk are shared and which are antibody-specific.

Our central finding is that the Fc glycans interact with numerous residues far from the N-glycosylation site, including residues on the Fab arms, in both antibodies. The allosteric networks sourced on the glycans point the same way, reaching beyond the Fc and into the Fabs, where they engage framework residues rather than the CDRs. Since framework residues shape the conformation of the hypervariable loops, this suggests a route by which Fc glycosylation could modulate antigen or receptor binding at a distance, a hypothesis that is now testable. Both antibodies proved highly flexible, undergoing large conformational changes throughout the trajectories despite different starting shapes set in part by their different hinge lengths. In both, and irrespective of glycosylation states, native contacts were continuously lost and replaced by new ones, so the overall structure remained stable throughout. This shared behaviour, rigid modules tethered by a flexible hinge, is the mechanical background against which the glycan effects were assessed.

Some opening states of CH2 domain are sampled, but the characterization is quite difficult depending on the criterion chosen. However, we do see a flexibility of the Fc domain, and glycosylation seems to have a modest impact on their dynamics.

Because each isotype is represented by a single molecule, and because the two mAbs differ at once in species, sequence, architecture, hinge length and glycan composition, these results are antibody-specific. They should not be generalised to other IgG2a or IgG4 monoclonal antibodies. Within that limit, the long-range contacts we describe reach toward the paratope and could influence the antigen binding. This study highlights the importance of considering full-length structures and the diversity of IgG scaffolds in glyco-engineering rather than the Fc alone.

## Data and Software Availability

The input files required to reproduce the simulations including starting coordinates (PDB/GRO), topology files, force-field parameters for the glycans, equilibration, production input files (.mdp), and the in-house analysis scripts (Python) and the full MD trajectories generated and analyzed during this study are available on Zenodo (https://zenodo.org/records/19680340). All software used in this work is open source or freely available for academic use: GROMACS (https://www.gromacs.org), CHARMM-GUI (https://www.charmm-gui.org), VMD (https://www.ks.uiuc.edu/Research/vmd/), and MDAnalysis (https://www.mdanalysis.org).

## Competing Interests

The authors declare no other competing interests.

## Author Contributions

Conceptualization, C.E, J.D, A.B and R.LFL; Methodology, C.E, J.D and R.LFL; Formal analysis, R. LFL & C.E.; Investigation, R. LFL; Data curation, R. LFL and A.B; Writing – original draft, R. LFL; Writing – review and editing, C.E, J.D and R.LFL; Visualization, R.LFL; Supervision, C.E, and J.D; Funding acquisition, ANRT. All authors read and approved the final manuscript.

## Acknowledgment

This work was supported by the French National Research and Technology Association (ANRT) through the CIFRE grant No. 2023-0969.

This project was also provided with computing HPC and storage resources by CINES thanks to the grant A20, number A0200711501 on the supercomputer ADASTRA and to which we are grateful for providing all the computational resources required for the MD simulations.

## Notes

### Competing Interest Statement

The authors have declared no competing interest.

### Summary of Updates

update supp mat with updated informations in the supp mat

