## Supplementary materials for "Molecular Dynamics Exploration of Two Full-Length IgG Antibodies: Impact of Glycosylation"

***Supplementary Methods.***

*Conformational and Dynamics Analysis.*

Most of the analyses were performed with the GROMACS 2023.3 tool suite[1] in particular the *rms* tool for the Root Mean Square Deviation (RMSD) and the *rmsf* tool for the Root Mean Square Fluctuation (RMSF) (averaged by residue). The RMSD was computed on the Cα atoms both globally (over the whole mAb) and locally (over Fab-1, Fab-2, Fc-1, and Fc-2 individually), with each domain defined in Table 1. In every case, the crystallographic structure of the corresponding antibody was used as the reference for alignment. The analyses were performed for each replicate but also on the whole set of trajectories depending on the measures considered. The mean and standard deviation across the three replicates were reported. The metrics were computed per replicates and were not pooled.

*Blockwise assessment of intra-replica stationarity.*

We assessed for each angular descriptor how much its distribution varies between consecutive time blocks within one replicate and compared internal variability with the glycosylated and aglycosylated differences measured on the same replicate.

For each replicate, only the conformations saved after ≥ 200 ns were kept. This gives N angular series, one per combination of descriptor (11), replicate (3), glycosylation state (2) and antibody (2). The integrated autocorrelation time of each series was estimated as detailed in the Statistical Analysis section of the main text. The circular descriptors φFab-1, φFab-2 and the hinge dihedral were expressed as their signed deviation from the circular mean to avoid the discontinuity at the 0–360° wrap-around.

Each series was then split into consecutive 100 ns blocks. We used this block size as the reference because shorter blocks leave too few of them for the slowest descriptors. Two dissimilarities were then compared, both measured as 1 − OVL, where OVL is the overlap between two distributions and 1 − OVL therefore measures how much they differ. The first is the internal dissimilarity: how much the distribution of a descriptor changes from one time block to the next within a single trajectory. The second is the condition dissimilarity: how much the glycosylated and aglycosylated series differ within the same replicate. If the internal dissimilarity is as large as the condition one, the difference between the two states cannot be told apart from the drift of a single trajectory over time.

Convergence of the standard error was checked separately with the Flyvbjerg–Petersen block-averaging procedure (Fig. S3).

*Dynamic Cross Correlation Matrix (DCCM).*

The DCCM measures the correlation of the motion of each residue with every other residue along the trajectory. We used the Pearson correlation, computed from the mean displacement vector of each residue:

$$\Delta r_i(t) = r_i(t) - \overline{r_i} \quad \text{(Supp. Eq. 1)}$$

where $r_i(t)$ is the position of residue i at time t and $\langle r_i \rangle$ is its mean position over the trajectory. The correlation between residues i and j is then:

$$C_{ij} = \frac{\langle \mathbf{\Delta r}_i \cdot \mathbf{\Delta r}_j \rangle}{\sqrt{\langle \mathbf{\Delta r}_i^2 \rangle \langle \mathbf{\Delta r}_j^2 \rangle}} \quad \text{(Supp. Eq.2)}$$

Before computing displacement vectors, all frames were aligned on the crystallographic structure of the corresponding antibody using the Cα atoms as the reference selection. The matrices were computed with the Bio3D library[2] implemented in R. The DCCM analysis was realized once the conformational convergence was reached by the mAbs at 200ns. It was computed per replicas and also with the concatenated simulations.

*Principal Component Analysis (PCA).*

We first selected the residues whose correlation or anticorrelation values differed most between the glycosylated and aglycosylated states, based on the difference-DCCM (glycosylated minus aglycosylated) computed for each mAb. Pairs of residues showing an absolute DCCM difference greater than 0.7 were retained. Because the number of residues impacted by glycans differs between the two mAbs, so does the number of retained pairs. For each selected pair, the Cα–Cα distance was then computed on the three replicates, after the 200ns equilibration time. The resulting distance ensembles were analysed by PCA using the MDAnalysis, NumPy, and Scikit-learn Python packages. This was computed using the pooled simulations.

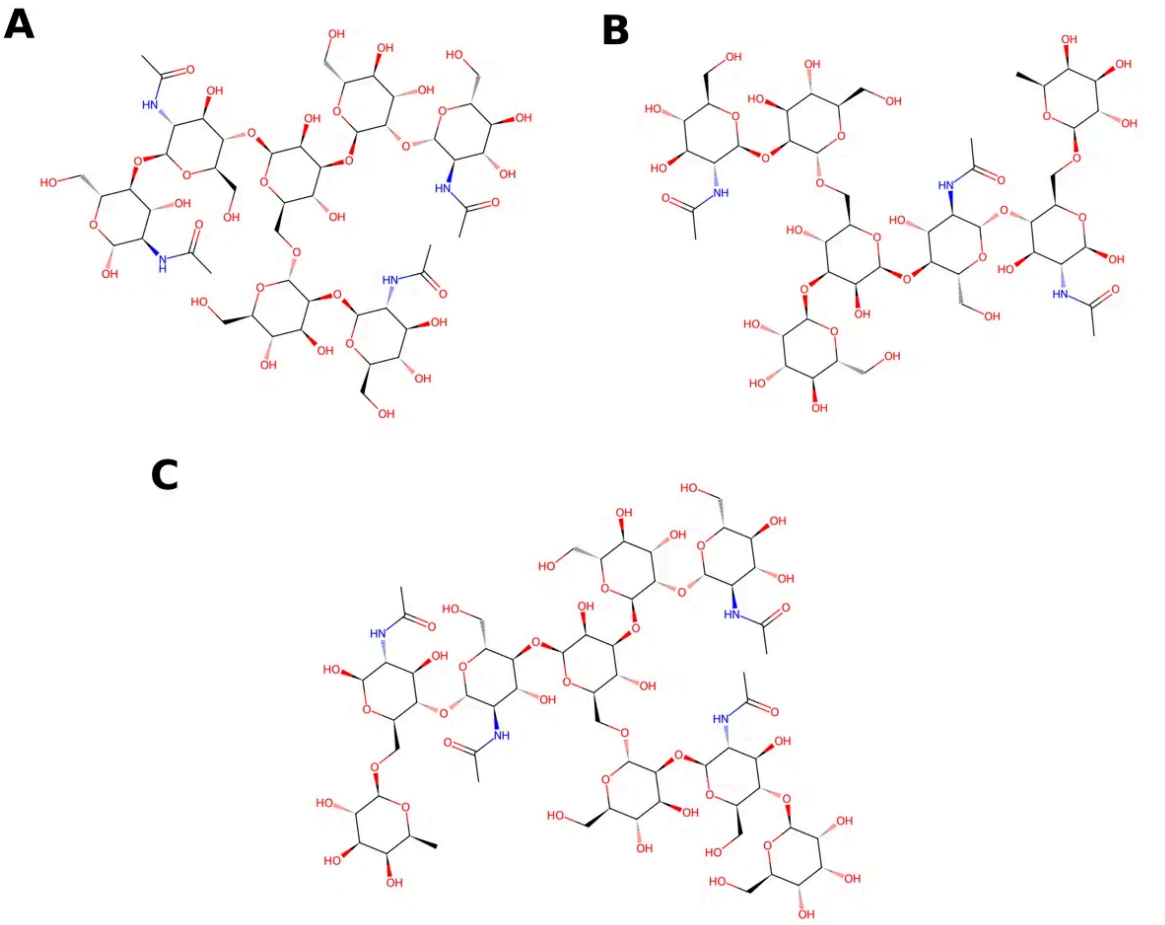


Figure S1 : CRAM representation of the glycans: CARA (A), CARB (B), and CARC_1 and CARC_2 (C). CARC_1 and CARC_2 are chemically identical and are shown together in panel C.

| | Identities | Positives |
|---|---|---|
| **Heavy Chains** | 59% | 75% |
| **Light Chains** | 62% | 76% |

Table S1 : Metrics from the sequence alignment between the heavy chains and between the light chains of PMB and Mab231, computed with the Needleman–Wunsch algorithm (NCBI). Identities and positives are reported as percentages.

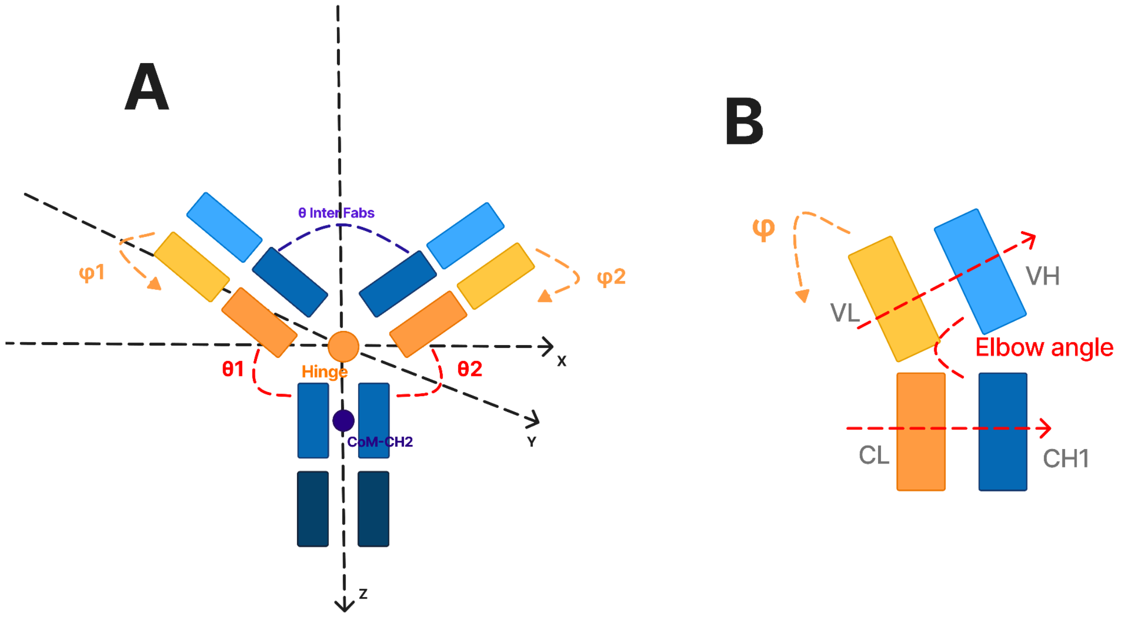


Figure S2 : Definition of the angular descriptors used to characterise antibody conformational states. (A) Coordinate system used to define Fab orientation relative to the Fc. The origin is the centre of mass of the hinge (orange dot). The z-axis runs from the hinge to the centre of mass of the CH2 domains, toward the Fc C-terminus; the x-axis lies in the antibody plane, perpendicular to z; the y-axis completes a right-handed frame. The polar angles $\theta_1$ and $\theta_2$ (red arcs) are the angles between the z-axis and the principal inertia axis of Fab1 and Fab2. The azimuthal angles $\varphi_1$ and $\varphi_2$ (green arrows) describe the rotation of each Fab arm around the Fc long axis. The inter-Fab angle θ_InterFabs (blue arc) is the angle between the two Fab inertia axes. Heavy-chain domains (VH, CH1, CH2, CH3) are in shades of blue, light-chain domains (VL, CL) in shades of green. (B) Elbow angle of a single Fab (red dashed lines), defined as the angle between the VH→VL and CH1→CL centre-of-mass vectors, measuring the bending between the variable and constant regions. The azimuthal angle φ (green arrow) is shown for reference.

***Supplementary Results.***

*Stability of a human IgG4 and a murine IgG2a with and without N-glycosylations.*

Both antibodies departed substantially from their starting conformation over the course of the simulations. After the initial rise, the global backbone RMSD of the full-length antibody plateaued at roughly 23 to 24 Å for aglycosylated PMB and 17 to 18 Å for glycosylated PMB, and at roughly 25 to 29 Å for aglycosylated Mab231 and 22 to 26 Å for glycosylated Mab231 (Fig. S4), the glycosylated form reaching lower global values than the aglycosylated one in both antibodies. Computed domain by domain, the RMSD of Fab-1, Fab-2 and Fc-2 stays low and flat throughout, between about 2 and 5 Å, so the global increase arises almost entirely from the change in the relative positioning of the Fc and Fab domains rather than from internal rearrangement. The only exception is Fc-1 of PMB, whose RMSD reached about 7 to 8 Å in both glycosylation states, indicating some internal deformation of this domain alone, possibly due to the loops in this region. Saporiti et al.[3] reported the same pattern for an IgG1, in which the local domains converge quickly while the global structure keeps rearranging, and attributed it to the high flexibility of monoclonal antibodies, whose structural equilibrium resides in the oscillation of the hinge and of the Fab arms rather than in a single well-defined global minimum.

The individual domains therefore reach a stable ensemble rapidly. The large-scale motions, by contrast, are the relative reorientations of the Fc and the two Fab arms about the hinge, and these have not reached a stationary state within a single replicate, because they are slow: the blockwise analysis of the angular descriptors (Fig. S3) gives integrated autocorrelation times with a median of 36.9 ns, 40 % of the series above 50 ns and the slowest reaching 200ns, so the slowest coordinates provide only four to ten independent samples over the roughly 800 ns retained per replicate. As previously observed for other full-length immunoglobulins, the main degrees of freedom of these large systems are the torsion angles of the hinge, located close to the glycosylation sites.

A single replicate therefore does not sample the full inter-domain landscape. The three replicates, however, explore potentially different regions of that landscape, so together they provide far from negligible sampling of the accessible conformational space. It is within this sampled space that we compared the glycosylated and aglycosylated forms, and there the sugars show no clear effect on the relative motions of the domains, as detailed in the following sections[4].

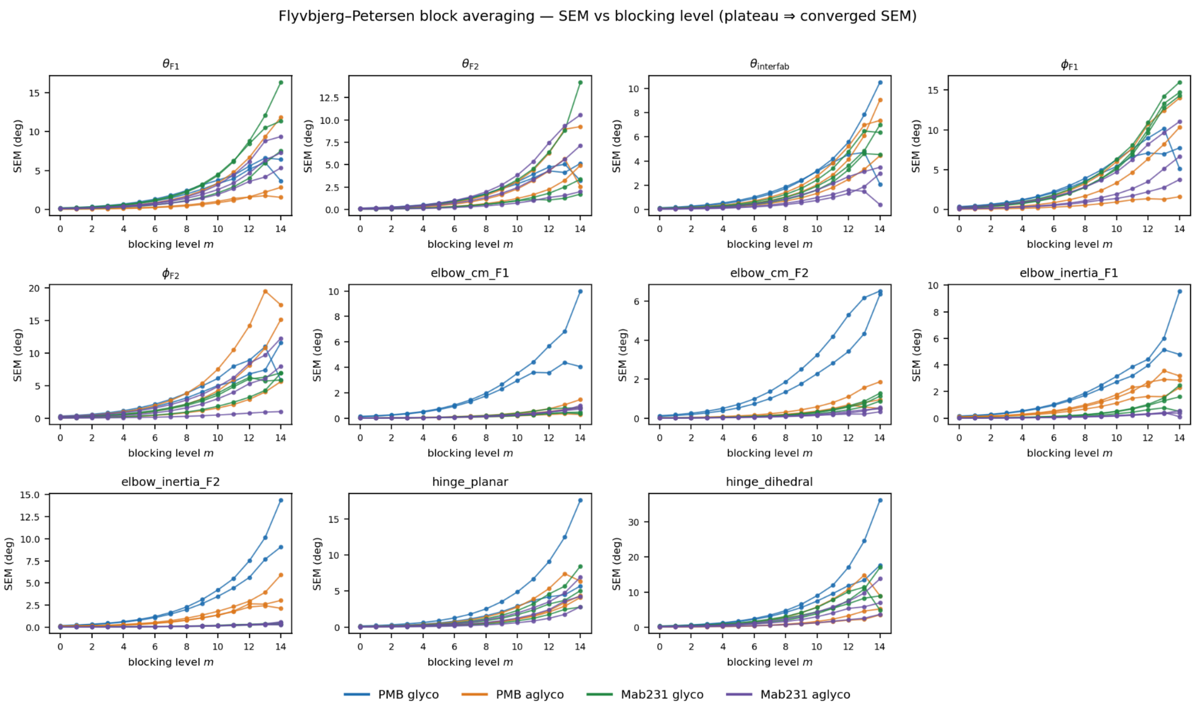


Fig S3 : Block averaged standard error of the angular descriptors, following Flyvbjerg and Petersen. The standard error of the mean is plotted against the blocking level m, each increment of m doubling the block length, with one panel per descriptor. Colours distinguish the antibody and the glycosylation state, and each line corresponds to one replicate. Series were analysed after removal of the first 200 ns ; the circular descriptors were treated on their signed deviation from the circular mean.

*RMSD Analysis.*

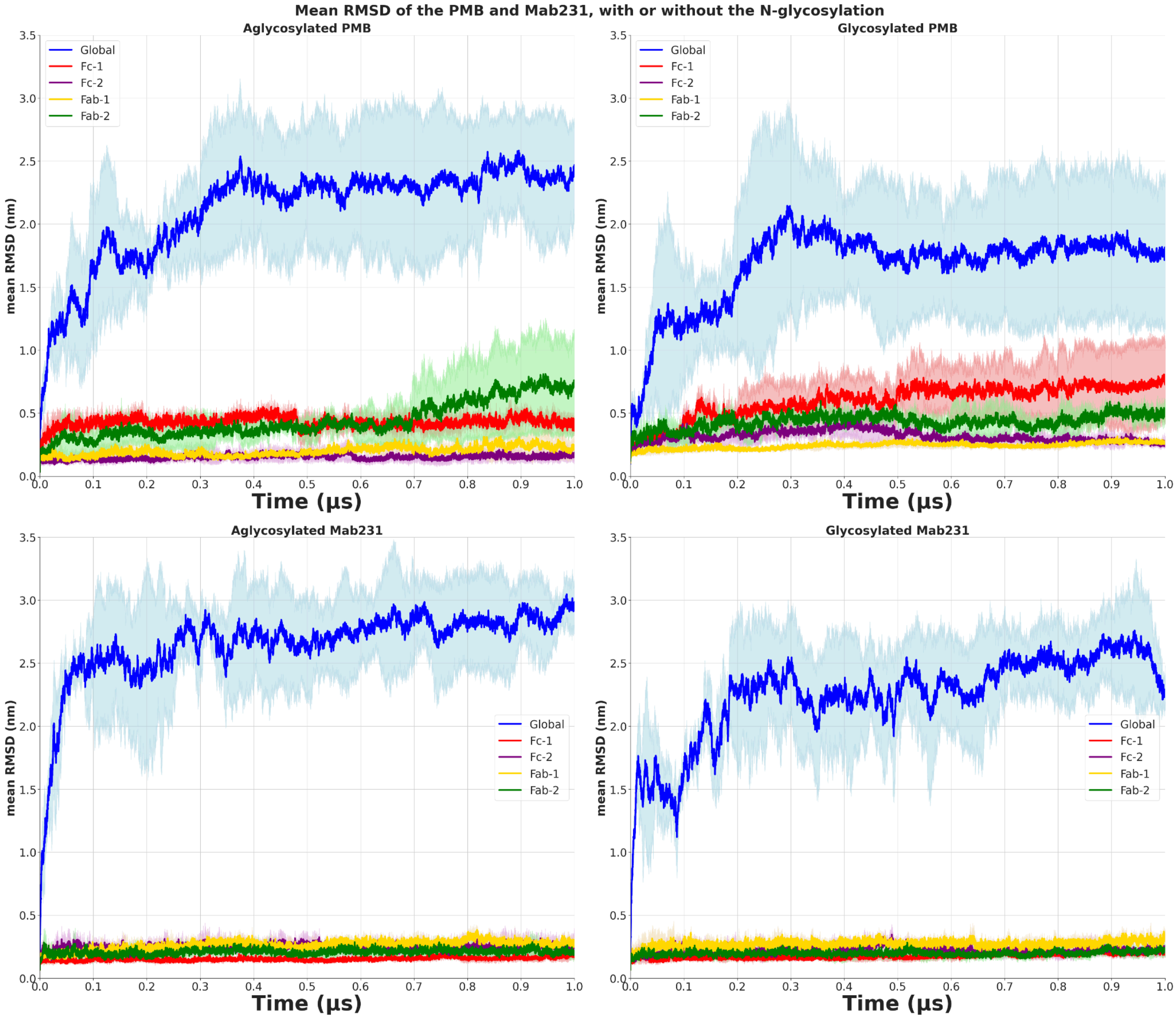


Figure S4 : Evolution of the mean RMSD on each replicate for PMB without (A) and with (B) the carbohydrates, and Mab231 without (C) and with (D) the N-glycosylation. The blue curve is the RMSD of the full backbone of the antibody, red is the Fc-1 chain, purple is the Fc-2 chain, green is the Fab-2 region and yellow is the Fab-1 region. Solid curves are the means across replicates and the shaded bands represent the standard-deviation values.

*Evolution of the average Native Contacts (NC) and Non-Natives contacts.*

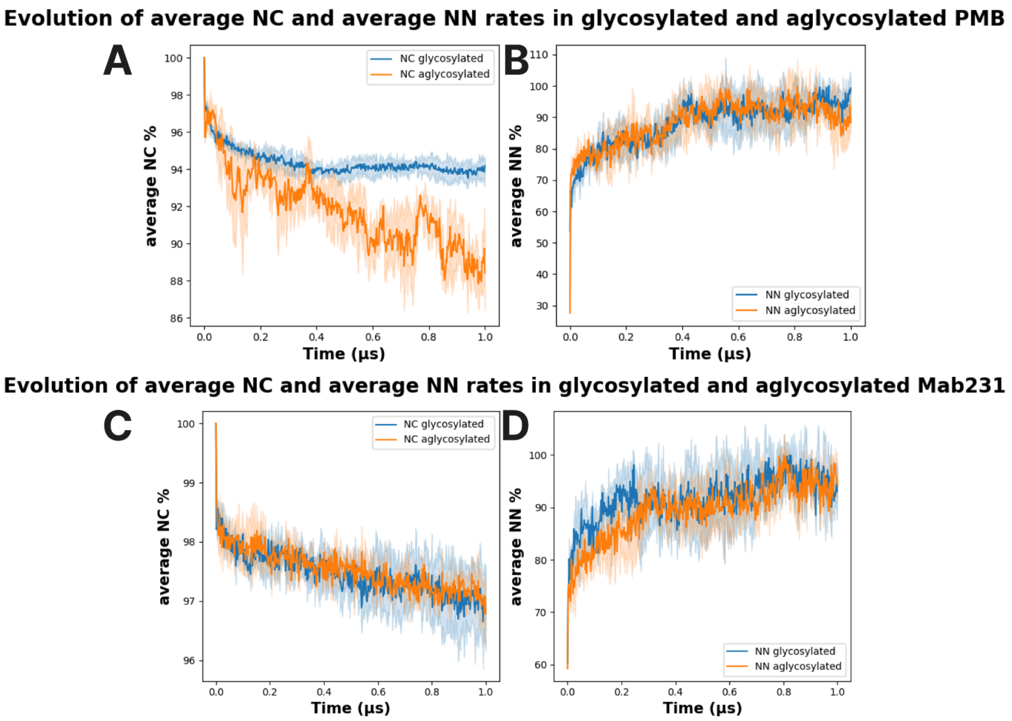


Figure S5 : Evolution of the rate of the average native contacts (NC, A & C) and non-native contacts (NN, B & D). Glycosylated systems are shown in blue and aglycosylated systems in orange in each panel. Values are shown for both PMB (A & B) and Mab231 (C & D). Solid curves are the means across replicates and the shaded bands represent the standard-deviation values.

*Flexibility of PMB and Mab231.*

Both antibodies exhibit large average fluctuation profiles (Fig. S6). , and the mean profiles differ between the glycosylated and the aglycosylated states,states but : considering the standard deviation between replicates and between the two forms that overlap, no significant effect of the glycosylation on the fluctuation profile can be established for PMB the glycosylated mean lies above the glycosylated one across most of the sequence, while for Mab231 the two means cross, the glycosylated form fluctuating more in Fab-1 and less in Fab-2.

This is consistent with the fact that the fluctuations of a full-length IgG are dominated by the reorientation of the arms around the hinge, which is precisely the motion that our replicates sample only partially.

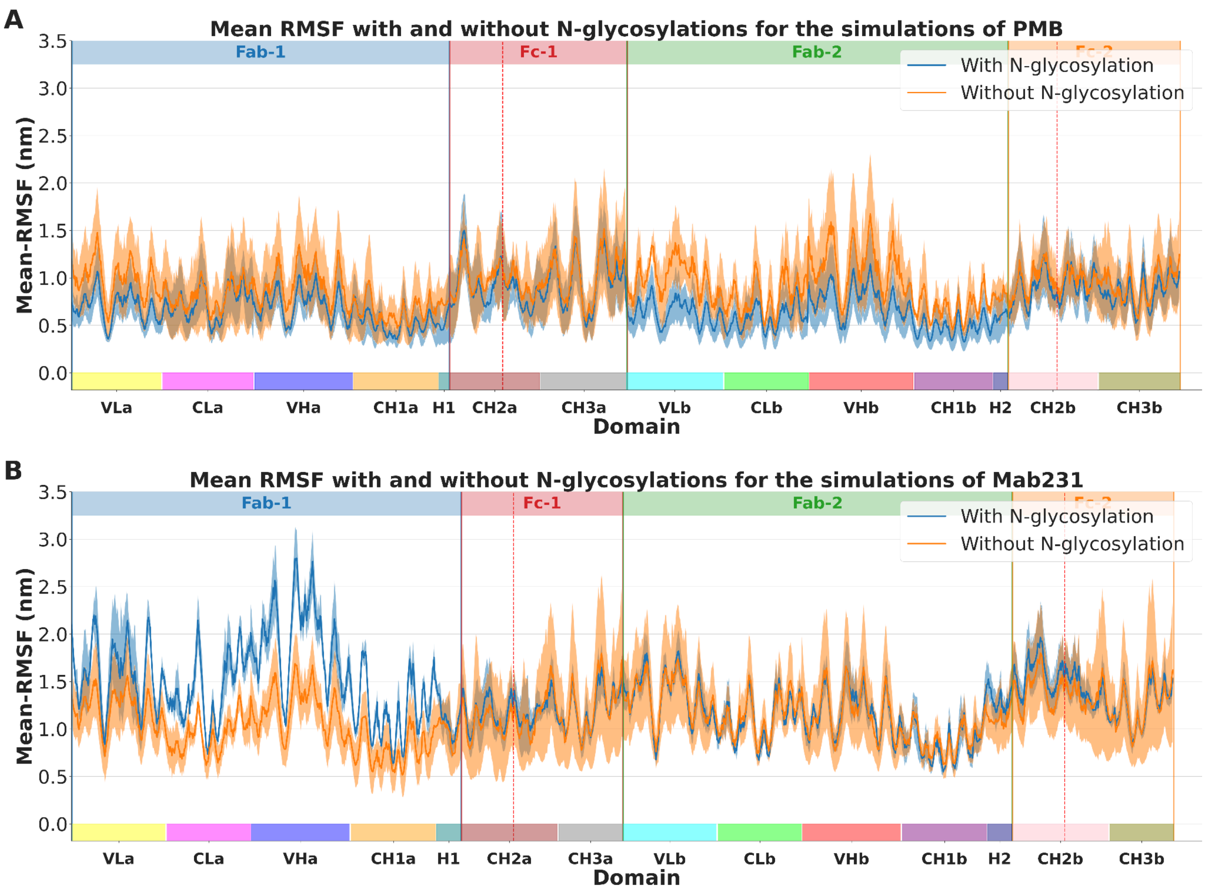


Figure S6 : Mean RMSF profiles for PMB (A) and Mab231 (B). The orange curve is the mean RMSF without N-glycosylation and the blue curve with N-glycosylation. Each red vertical line marks the position of the residue linked to the carbohydrate. A dashed vertical line separates two domains that are disconnected in the structure. Domain boundaries are indicated along the x-axis (Fab-1, Fc-1, Fab-2, Fc-2, with their VL/CL/VH/CH1/CH2/CH3 subdomains and hinges H1/H2). Solid curves are the means across replicates and the shaded bands represent the standard-deviation values.

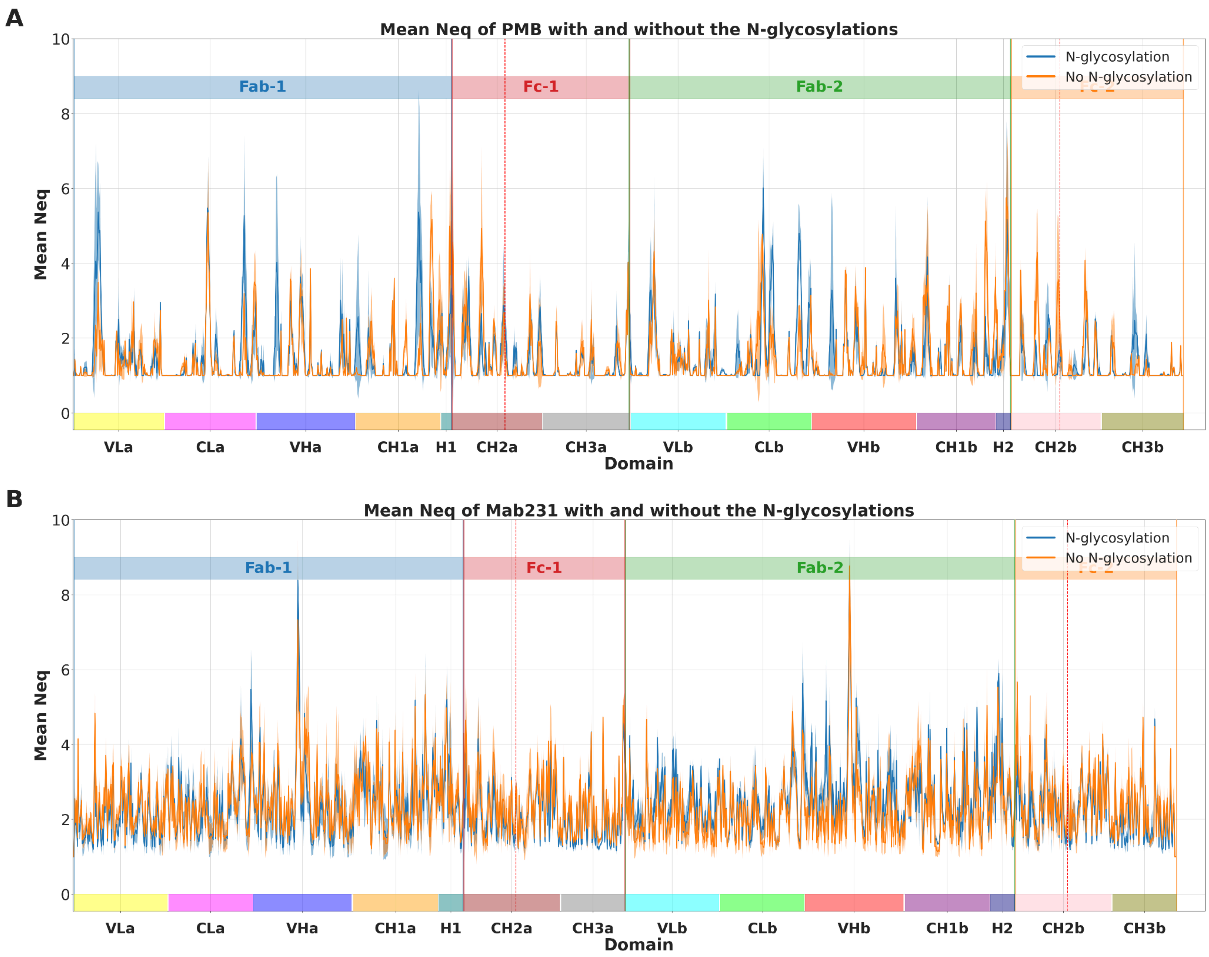


Figure S7 : Mean Neq for PMB (A) and Mab231 (B) in both conditions. The orange curve is the mean Neq without N-glycosylation and the blue curve with N-glycosylation. Each red vertical line marks the position of the residue linked to the carbohydrate. A dashed vertical line separates two domains that are disconnected in the structure. Domain boundaries are indicated along the x-axis. Solid curves are the means across replicates and the shaded bands represent the standard-deviation values.

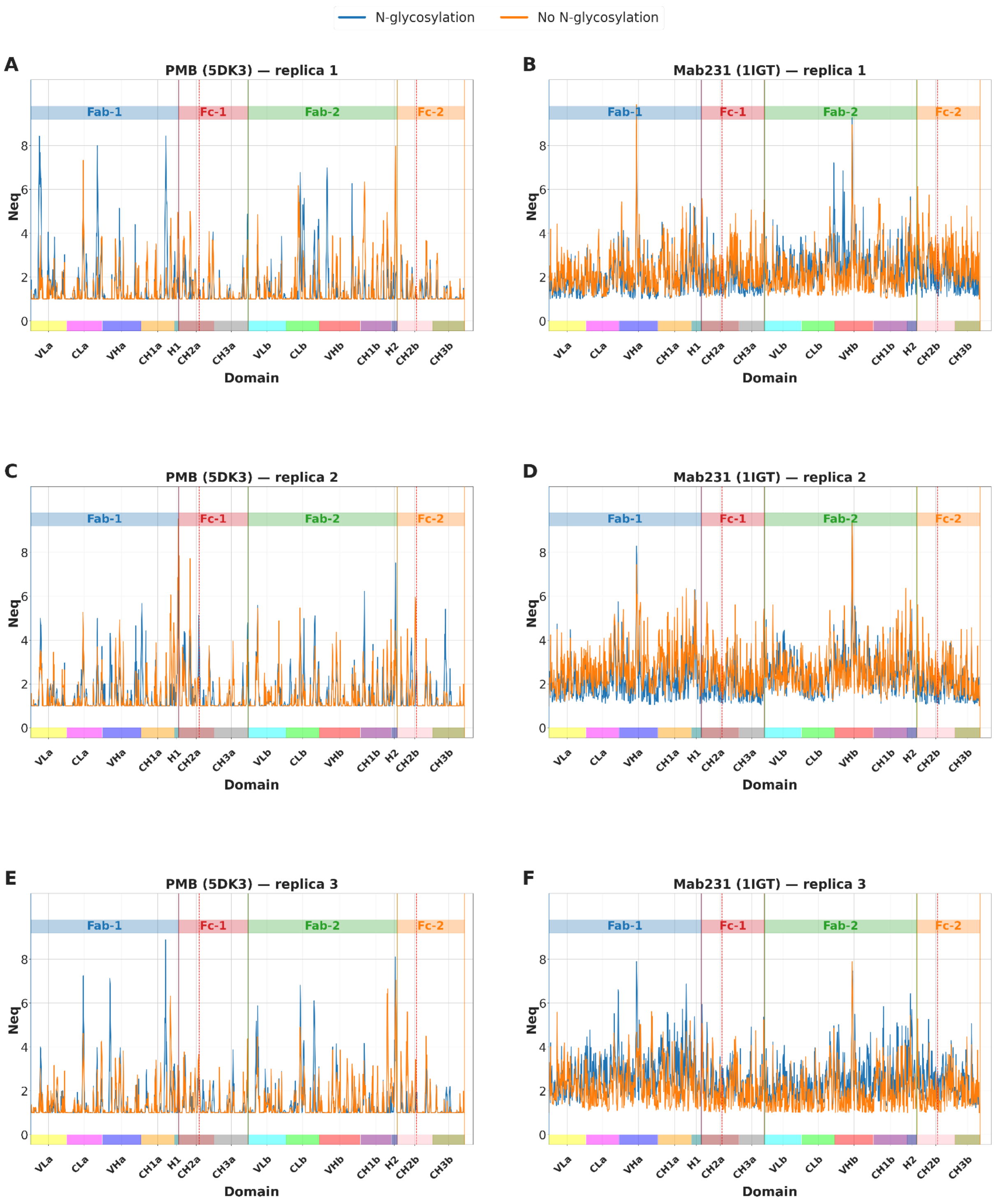


Figure S8 : Per-replicate mean Neq for PMB (5DK3, A/C/E) and Mab231 (1IGT, B/D/F), for replicas 1, 2 and 3 respectively. The orange curve is the mean Neq without N-glycosylation and the blue curve with N-glycosylation. Each red vertical line marks the position of the residue linked to the carbohydrate. Domain boundaries (Fab-1, Fc-1, Fab-2, Fc-2 and their subdomains) are indicated along the x-axis.

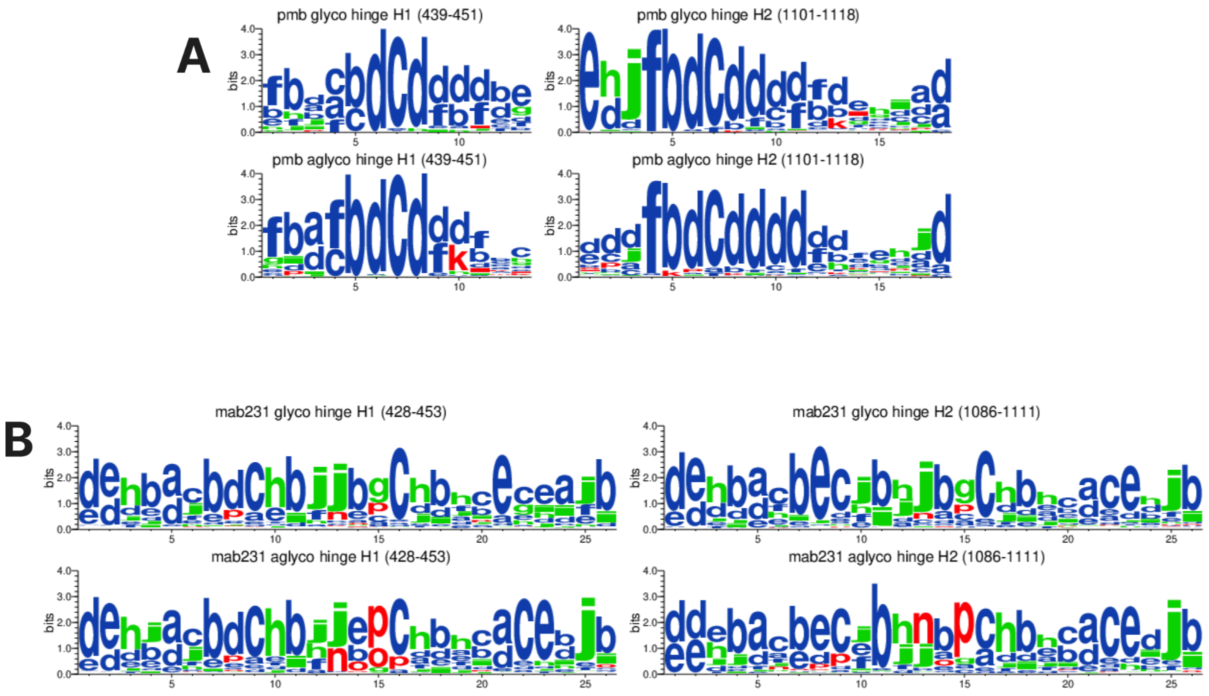


Figure S9 : WebLogo of the hinge domains of PMB (A) and Mab231 (B), with and without the N-glycosylations. For each antibody the two hinge regions (H1 and H2) are shown for the glycosylated and aglycosylated states; residue ranges are indicated above each logo and letter height is expressed in bits.

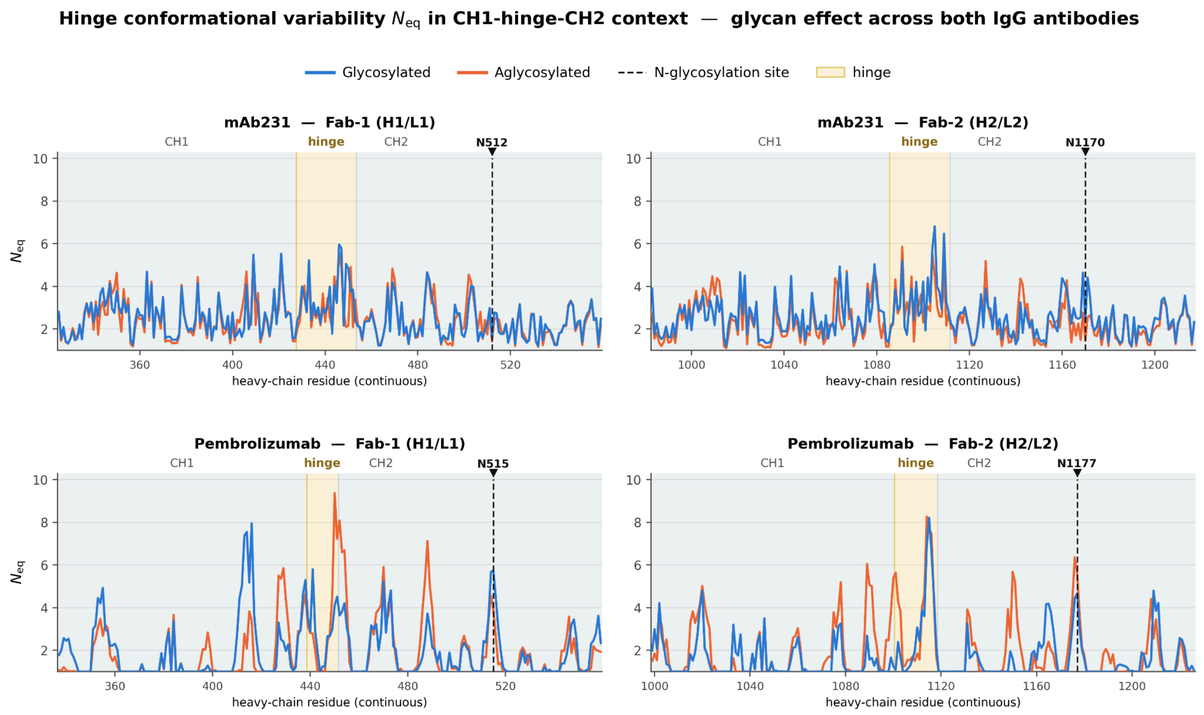


Figure S10 : Hinge conformational variability (Neq) in the CH1–hinge–CH2 context, comparing the average glycosylated (blue) and aglycosylated (orange) Neq for Mab231 and PMB, from the CH1 domain to the CH2 domain, for Fab-1 (H1/L1) and Fab-2 (H2/L2). The hinge region is highlighted in shading and the glycosylated Asn (N512/N1170 for Mab231; N515/N1177 for

PMB) is indicated by a black dashed line. The x-axis is the continuous heavy-chain residue numbering.

*Exploration of the cross talk between subdomains.*

We computed the dynamic cross correlation matrix (DCCM), which evaluates the correlated and anticorrelated motions between residues, with and without the sugars. The DCCMs computed per replicate show the same qualitative behaviour in each replicate, for both antibodies (Fig. S11 & S12 & S13). The absolute difference between the glycosylated and aglycosylated matrices (Fig. 5) contains many differing pairs, particularly for PMB, with the largest apparent changes between the Fc and the Fabs, for example CLb relative to CH2a and CH2a relative to CH1b.

The correlated motions (DCCM) and the orientation of the Fab arms relative to the Fc are examined below, are affected by this same reorientation. We therefore test each of them by checking that the result does not depend on the frame superposition used, and by verifying that the glycosylated-versus-aglycosylated difference exceeds the variability between replicates of the same state.

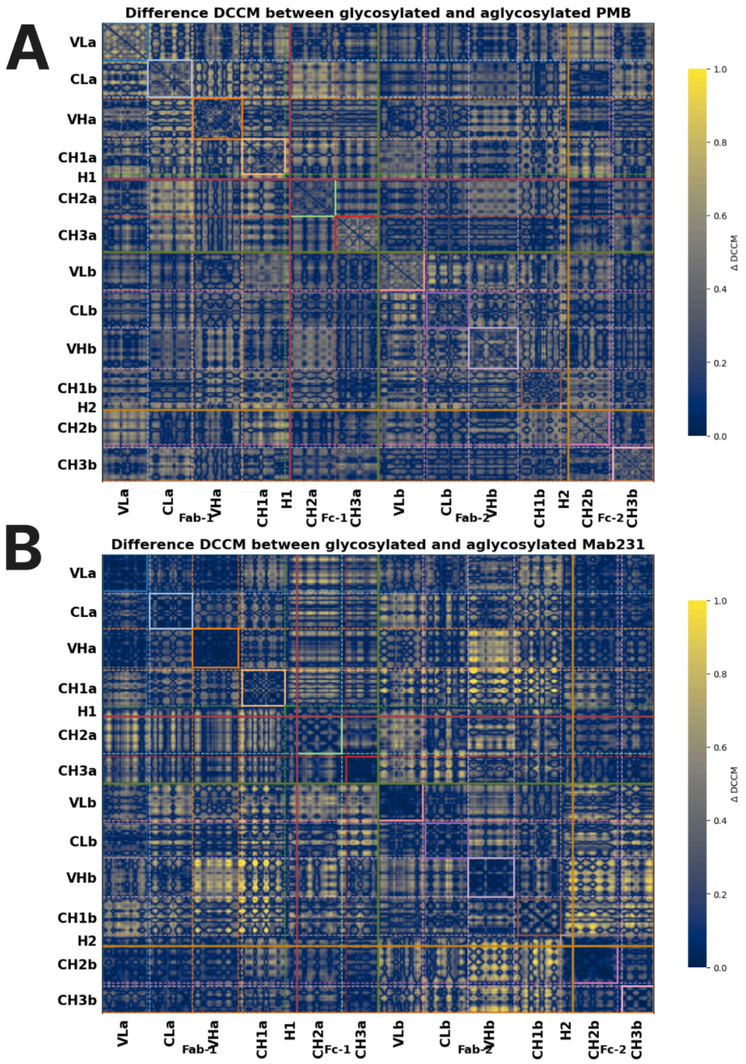


Figure S11: Difference DCCM computed from the glycosylated and aglycosylated DCCMs for PMB (A) and Mab231 (B). Values are absolute differences (glycosylated minus aglycosylated), with the colour bar (ΔDCCM) ranging from 0 to 1. The x- and y-axes list the residues grouped by subdomain (VL, CL, VH, CH1, H1, CH2, CH3 for each chain), and coloured frames along the axes delimit the Fab-1, Fc-1, Fab-2 and Fc-2 blocks.

*Does the protein adopt preferred conformations depending on the glycosylated state?*

The difference maps above identify the residue pairs whose cross-correlation differs most between the two states, and it is worth asking whether the distances between those pairs separate the two states in conformational space. We therefore retained the pairs for which the DCCM difference exceeds 0.7 and analysed the distribution of their distances by dimensionality reduction.

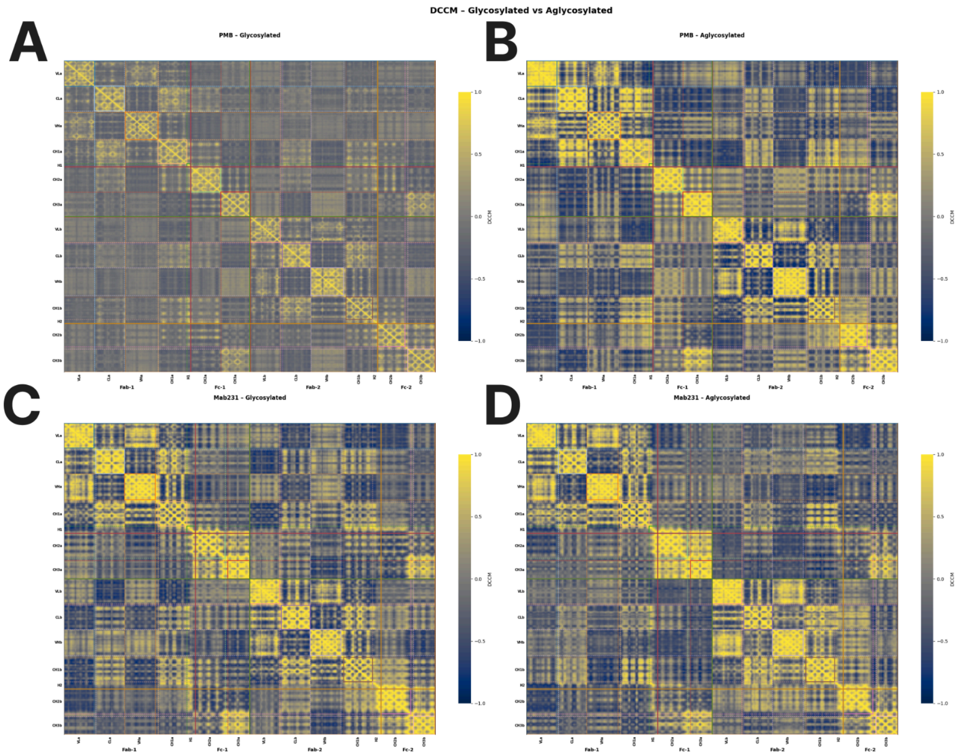


Figure S12: DCCM for PMB (A, B) and Mab231 (C, D), with (A, C) and without (B, D) the N-glycosylation. Values range from −1 to 1 (colour bar). The x- and y-axes represent the amino acids of the protein. The orange square delimits the amino acids of Fab-1, the green square those of Fab-2, the red square those of Fc-1 and the pink square those of Fc-2.

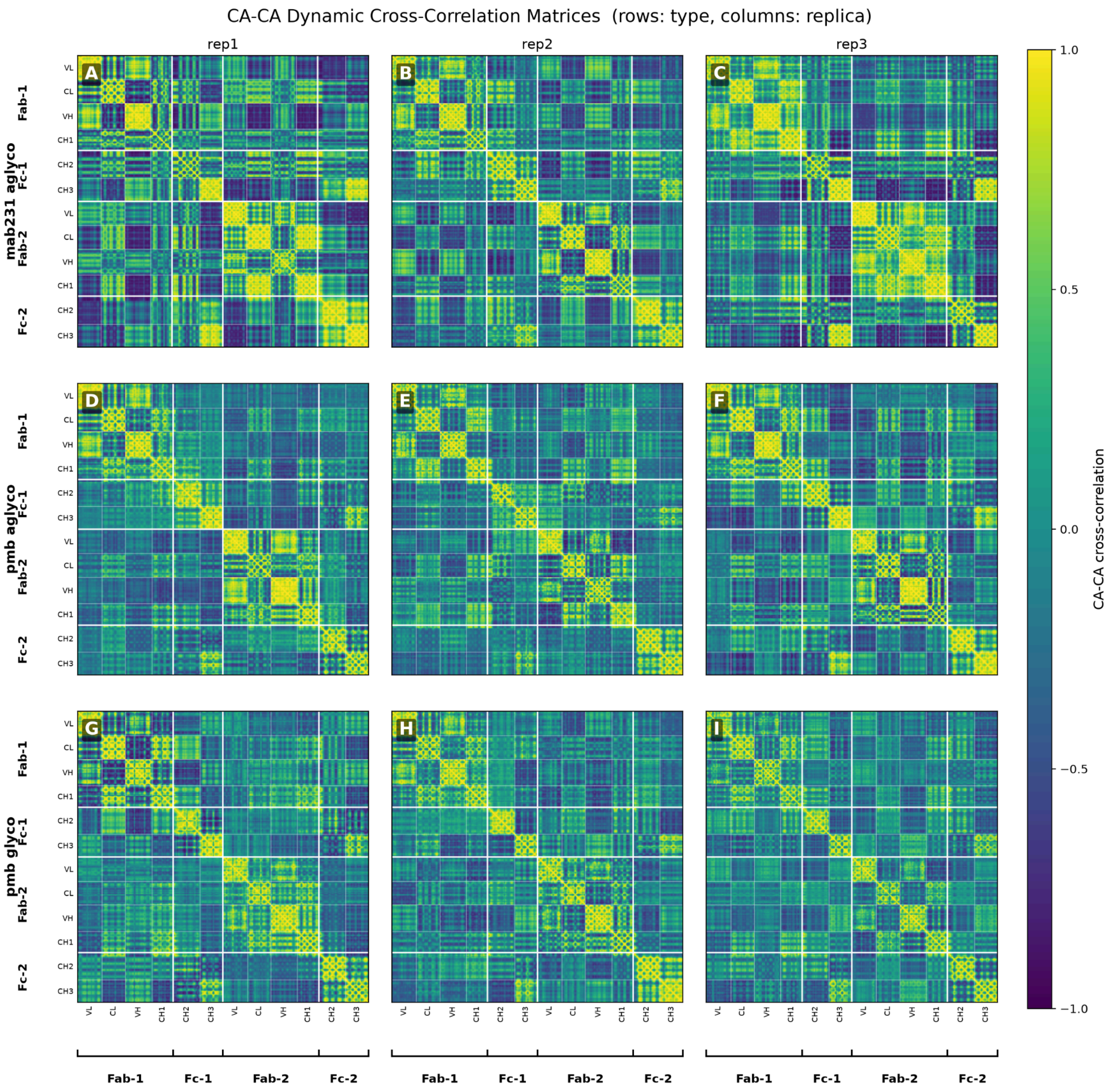


Figure S13: Per-replicate Cα–Cα DCCMs for Mab231 aglycosylated (A, B, C) and glycosylated (D, E, F), and PMB aglycosylated (G, H, I) and glycosylated (J, K, L); columns correspond to replicas 1, 2 and 3. The colour bar gives the Cα–Cα cross-correlation from −1 to 1. Subdomain blocks (A/B/C/D) are indicated along both axes.

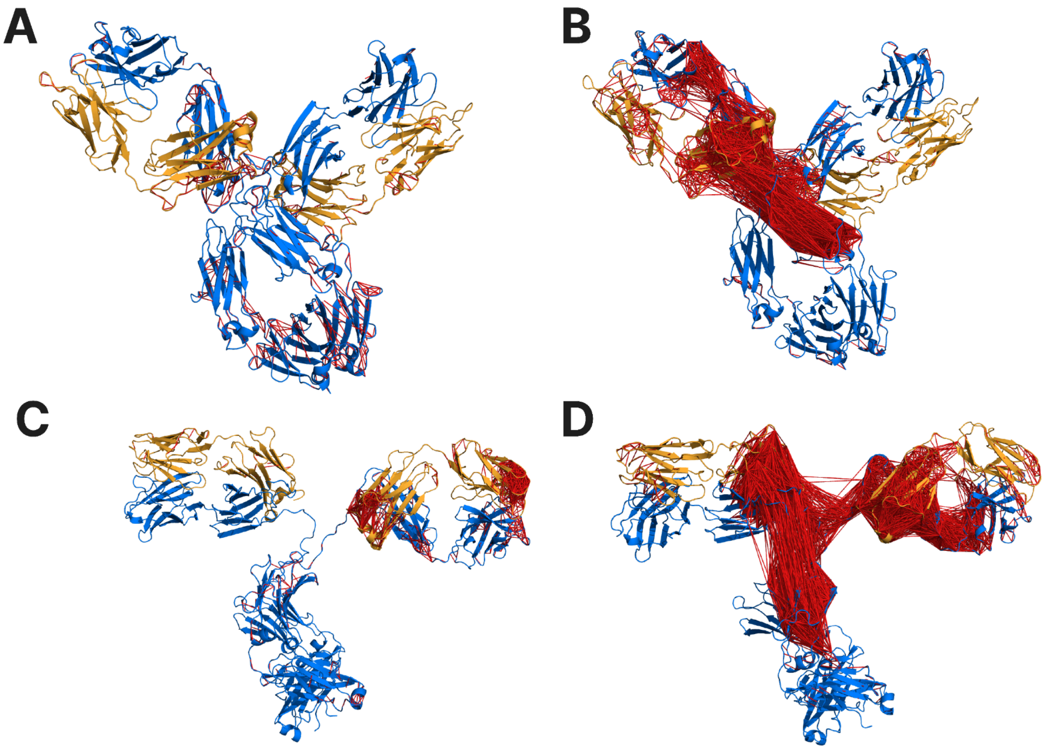


Figure S14: Projection on the structure of the amino acids which have correlated movements between each other. With and without the presence of the N-glycosylation. PMB is represented without the glycosylation in A and with B. Mab231 is represented without the glycosylation in C and with D. A red link exists when the correlation is between 0.9 and 1 between two amino acids.

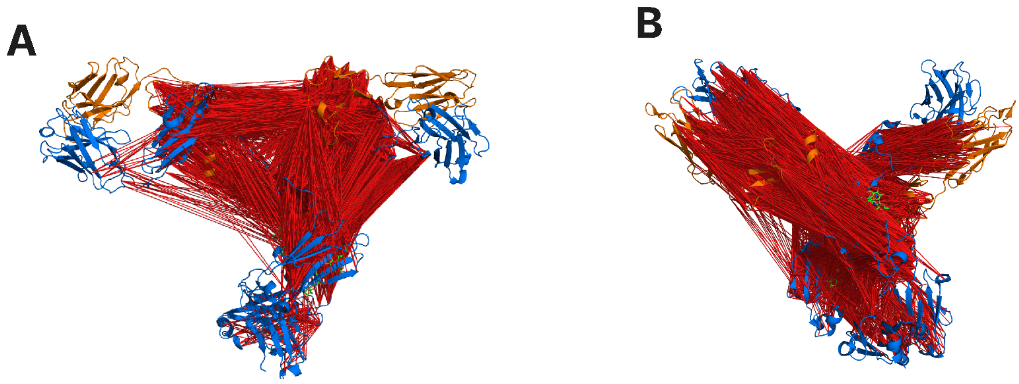


Figure S15: Projection of the pairs of residues that present the difference in the difference DCCM for Mab231 (A) and PMB (B).

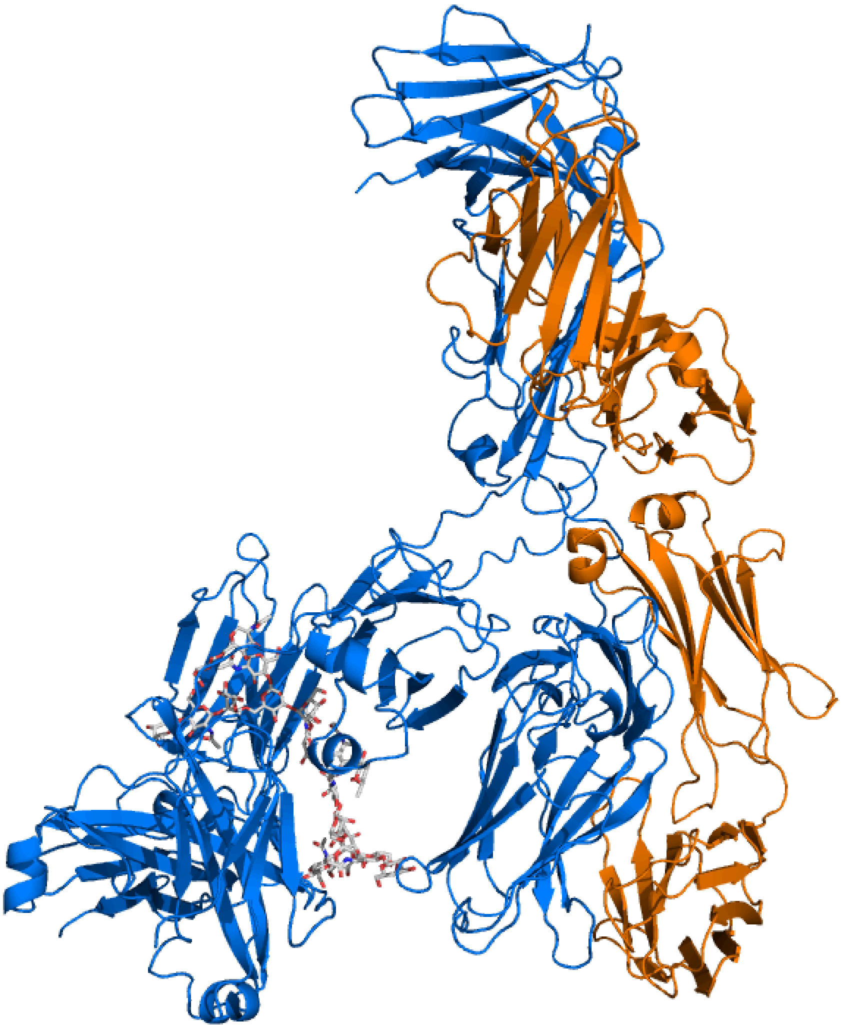

Figure S16: λ conformation obtained by Mab231 in the simulations.The blue chains are the heavy chains and the orange chains are the light chains. The glycans are represented in grey licorices.

*Reduction dimension methods to analyze glycosylated versus aglycosylated monoclonal antibodies.*

Figure S17 presents the projections of the conformations into the PC1 and 2 subspaces. For PMB (Fig. S11), 205,632 pairs of residues were retained. A large number that is associated with the different Fab-FC arm conformations sampled in all simulations. PC1 accounts for 28.2% of the variance and PC2 for 16.8%. First, there is no clear separation between the glycosylated and the aglycosylated states. Second, the conformations coming from different replicates of a given state are as distinct from one another as the two states are from each other.

For Mab231, 6,506 pairs were retained (Fig. S11.B), far fewer than for PMB, consistent with its two glycosylation states sampling more similar Fab-Fc configurations. PC1 accounts for 56.1% of the variance. The same conclusion holds: no clear distinction between the two glycosylation states emerges from the relative motions of the selected residue pairs. This conclusion does not depend on the threshold, as it remains unchanged when values other than 0.7 are used, other test values were also tried.

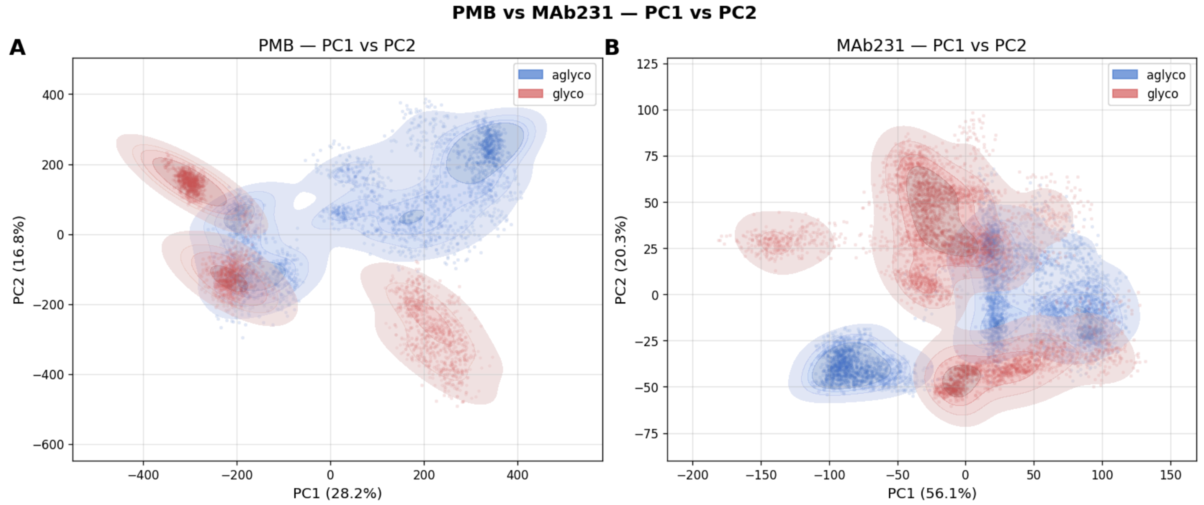


Figure S17: Projection onto the first two principal components (PC1, PC2) of the Cα–Cα distance ensembles for PMB (A) and Mab231 (B). Points are coloured by glycosylation state, glycosylated in red and aglycosylated in blue; the percentage of variance explained is indicated on each axis. Shaded contours indicate the density of each state.

*Angle analysis.*

To distinguish genuine glycan associated shifts from replicate to replicate variability, we examined each angular descriptor replicate by replicate (Fig. S18 & S19, summarised in Table S2 & S3). For PMB, the sign of the shift in θFab-1, θFab-2 and θInterFab is preserved in 3/3, 3/3 and 2/3 replicates respectively, with median per-replicate rank-biserial effect sizes of −0.65, −0.41 and −0.14, and overlap coefficients of 0.42, 0.60 and 0.59. For Mab231, the θFab-1 and θFab-2 shifts are consistent in 3/3 and 2/3 replicates, with median effect sizes of −0.38 and +0.34 and overlap coefficients of 0.58 and 0.40, whereas θInterFab shows a weaker and less reproducible effect (median r = +0.13, overlap 0.52, sign preserved in 2/3 replicates). The effect sizes and overlap coefficients quoted here and in Tables S2 and S3 are the medians of the three per-replicate values, not quantities computed on the pooled data. A consistent sign across replicates, however, is a necessary but not a sufficient condition for a resolved effect: a shift can be reproducible in direction while remaining smaller than the spread between replicates of the same state. We therefore applied to the angular descriptors the same two tests used above for the DCCM. Only θFab-1 keeps a consistent sign in all three replicates of both antibodies; θFab-2 does so for PMB only, and θinter-Fab for neither. On the noise bench, θFab-1 is the strongest of the three, with a signal-to-noise ratio near or slightly above one (0.94 for PMB, 1.24 for Mab231), yet even for θFab-1 the true glycosylated-versus-aglycosylated split is not the largest among the ten possible replicate relabellings.

Three further descriptors were not tested for reproducibility and are shown for illustration only (Fig. S18 & S19): the azimuthal Fab rotations φ1 and φ2, the elbow angle of each Fab, and the hinge dihedral. The elbow angles stay narrow around their canonical range in both antibodies and both states, consistent with the Fab modules behaving as rigid units, while the azimuthal rotations and the hinge dihedral are broad and multimodal, as expected for a full-length IgG whose arms reorient slowly around the hinge and sample the characteristic Y, T and λ shape.

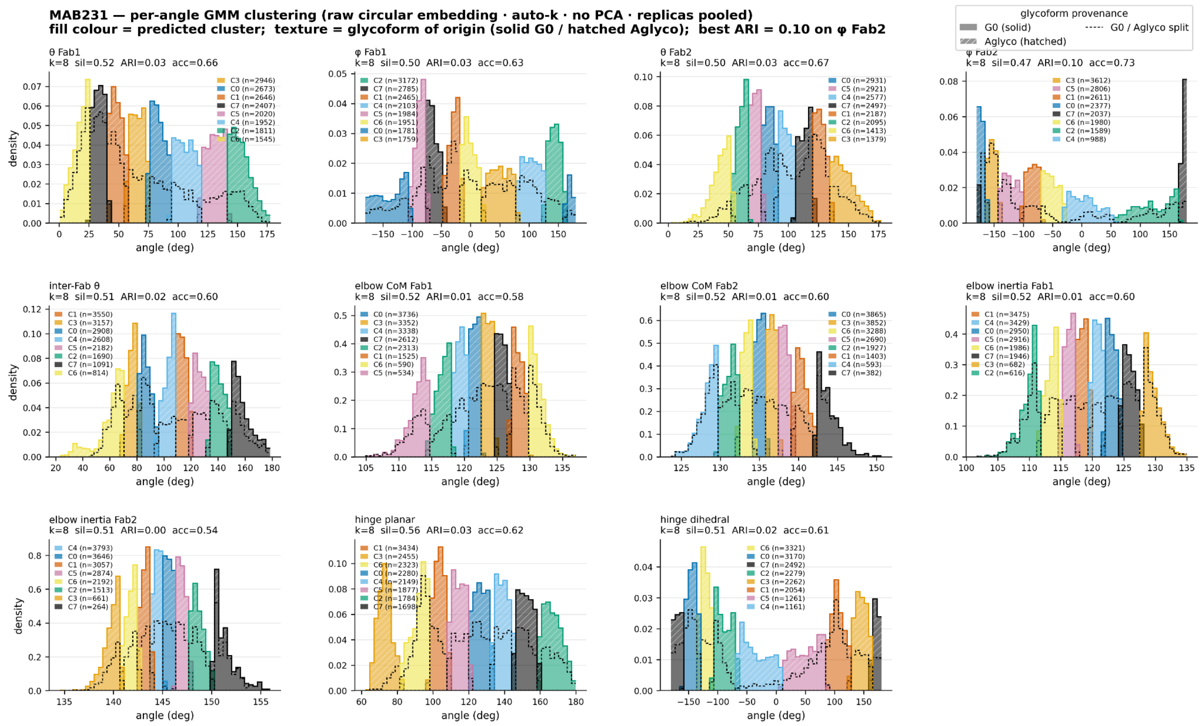


Figure S18: Per-angle GMM clustering of the angle values adopted by Mab231 (raw circular embedding, auto-k, no PCA, replicas pooled), for θFab-1, θFab-2, θInterFab, φFab-1, φFab-2, the elbow angles (centre-of-mass and inertia definitions of each Fab) and the hinge dihedral. Fill colour indicates the predicted cluster and texture indicates the glycoform of origin (solid = glycosylated G0, hatched = aglycosylated), the two glycoforms being separated by dashed black lines on the distributions. Cluster medoids are indicated by an arrow of the cluster colour, and the crystallographic structure by a solid red line. Per-panel silhouette, ARI and accuracy values, and per-cluster counts, are reported on each panel.

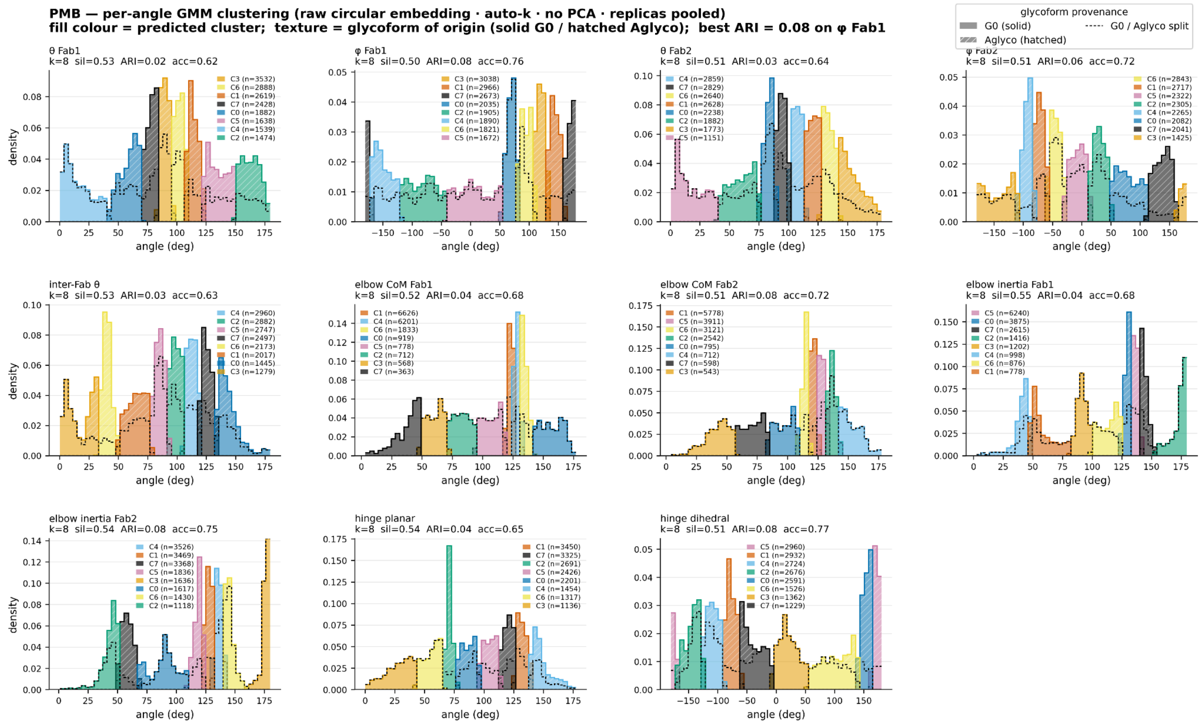


Figure S19: Per-angle GMM clustering of the angle values adopted by PMB (raw circular embedding, auto-k, no PCA, replicas pooled), for the same descriptors as in Figure S18. Fill colour indicates the predicted cluster and texture indicates the glycoform of origin (solid = glycosylated G0, hatched = aglycosylated), separated by dashed black lines. Cluster medoids are indicated by an arrow of the cluster colour and the crystallographic structure by a solid red line. Per-panel silhouette, ARI and accuracy values, and per-cluster counts, are reported on each panel.

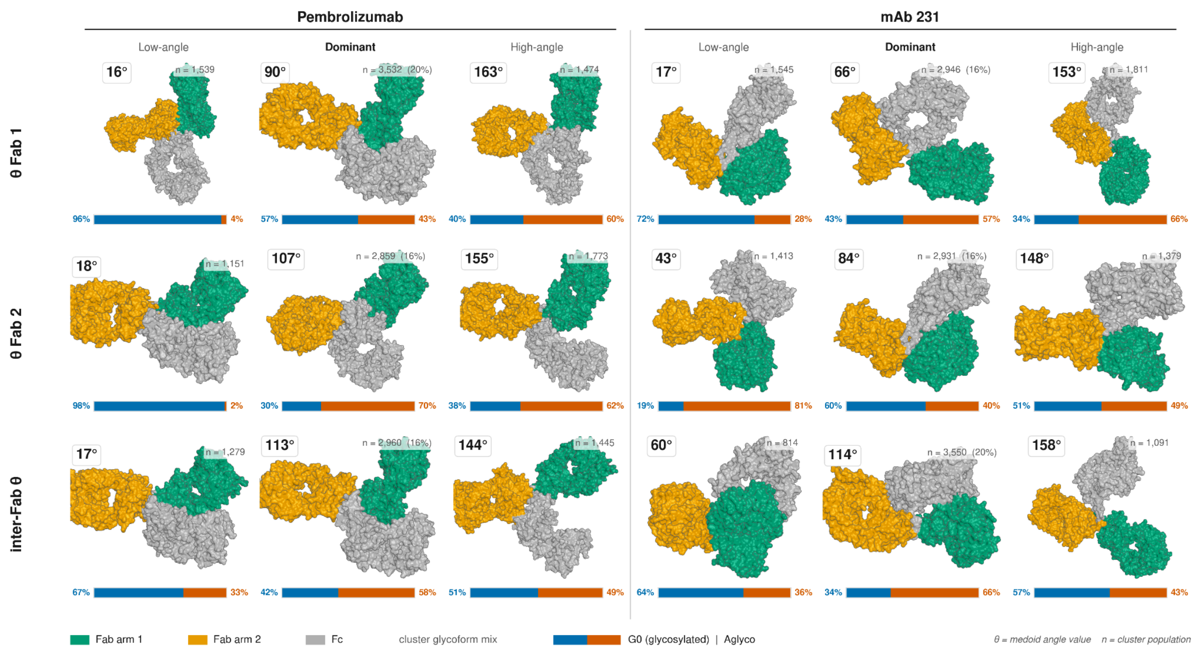


Figure S20 : Representative full-antibody conformations of the Fab-orientation clusters. For each angular distribution (θFab-1, θFab-2, θInter-Fab) the two extreme-angle clusters flanking the dominant (most populated) conformer are shown, using the GMM medoids (whole-antibody superposition), for PMB and Mab231. Fab arm 1 is green, Fab arm 2 is orange and the Fc is grey. The medoid angle value (θ) and cluster population (n) are indicated for each conformer, and the horizontal bar below each structure gives the glycoform mix of the cluster (G0 glycosylated vs aglycosylated).

| Angle | Cluster | Color | Pembrolizumab (IgG4) | | | |
|---|---|---|---|---|---|---|
| | | | Frames | Total % | Glyco % | Aglyco % |
| θ Fab-1 | C0 | | 1,882 | 10.46 | 55.2 | 44.8 |
| | C1 | | 2,619 | 14.55 | 31.1 | 68.9 |
| | C2 | | 1,474 | 8.19 | 40.2 | 59.8 |
| | C3 | | 3,532 | 19.62 | 57.0 | 43.0 |
| | C4 | | 1,539 | 8.55 | 95.7 | 4.3 |
| | C5 | | 1,638 | 9.10 | 43.5 | 56.5 |
| | C6 | | 2,888 | 16.04 | 51.4 | 48.6 |
| | C7 | | 2,428 | 13.49 | 35.8 | 64.2 |
| θ Fab-2 | C0 | | 2,238 | 12.43 | 69.8 | 30.3 |
| | C1 | | 2,628 | 14.60 | 40.6 | 59.4 |
| | C2 | | 1,882 | 10.46 | 65.8 | 34.2 |
| | C3 | | 1,773 | 9.85 | 38.0 | 62.0 |
| | C4 | | 2,859 | 15.88 | 29.7 | 70.3 |
| | C5 | | 1,151 | 6.39 | 98.3 | 1.7 |
| | C6 | | 2,640 | 14.67 | 42.1 | 57.9 |
| | C7 | | 2,829 | 15.72 | 48.4 | 51.6 |
| inter-Fab θ | C0 | | 1,445 | 8.03 | 51.3 | 48.7 |
| | C1 | | 2,017 | 11.21 | 57.8 | 42.2 |
| | C2 | | 2,882 | 16.01 | 54.1 | 45.9 |

| Angle | Cluster | Color | Pembrolizumab (IgG4) | | | |
|---|---|---|---|---|---|---|
| | | | Frames | Total % | Glyco % | Aglyco % |
| | C3 | | 1,279 | 7.11 | 67.3 | 32.7 |
| | C4 | | 2,960 | 16.44 | 42.4 | 57.6 |
| | C5 | | 2,747 | 15.26 | 74.6 | 25.4 |
| | C6 | | 2,173 | 12.07 | 16.3 | 83.7 |
| | C7 | | 2,497 | 13.87 | 40.6 | 59.4 |

| Angle | Cluster | Color | Frames | Mab231 (IgG2a) | | |
|---|---|---|---|---|---|---|
| | | | | Total % | Glyco % | Aglyco % |
| θ Fab-1 | C0 | | 2,673 | 14.85 | 39.2 | 60.8 |
| | C1 | | 2,646 | 14.70 | 63.8 | 36.2 |
| | C2 | | 1,811 | 10.06 | 33.6 | 66.4 |
| | C3 | | 2,946 | 16.37 | 42.9 | 57.1 |
| | C4 | | 1,952 | 10.84 | 38.5 | 61.5 |
| | C5 | | 2,020 | 11.22 | 30.1 | 69.9 |
| | C6 | | 1,545 | 8.58 | 72.5 | 27.5 |
| | C7 | | 2,407 | 13.37 | 79.5 | 20.5 |
| θ Fab-2 | C0 | | 2,931 | 16.28 | 60.0 | 40.0 |
| | C1 | | 2,187 | 12.15 | 74.2 | 25.8 |
| | C2 | | 2,095 | 11.64 | 28.4 | 71.7 |
| | C3 | | 1,379 | 7.66 | 50.6 | 49.4 |
| | C4 | | 2,577 | 14.32 | 59.7 | 40.3 |
| | C5 | | 2,921 | 16.23 | 29.4 | 70.6 |
| | C6 | | 1,413 | 7.85 | 19.3 | 80.8 |
| | C7 | | 2,497 | 13.87 | 66.4 | 33.6 |
| inter-Fab θ | C0 | | 2,908 | 16.16 | 70.3 | 29.7 |
| | C1 | | 3,550 | 19.72 | 33.5 | 66.5 |
| | C2 | | 1,690 | 9.39 | 58.1 | 41.9 |
| | C3 | | 3,157 | 17.54 | 48.8 | 51.2 |
| | C4 | | 2,608 | 14.49 | 39.5 | 60.5 |
| | C5 | | 2,182 | 12.12 | 48.9 | 51.1 |
| | C6 | | 814 | 4.52 | 64.3 | 35.8 |
| | C7 | | 1,091 | 6.06 | 57.1 | 42.9 |

Table S2 : Cluster populations of the Fab orientation angles forPMB and Mab231. For each angular descriptor (θ Fab-1, θ Fab-2 and the inter-Fab angle θ), the conformations were partitioned into eight clusters (C0–C7) using Gaussian Mixture Model clustering (k = 8, selected by the Bayesian Information Criterion).

*Interactions between glycans and monoclonal antibodies.*

**A**

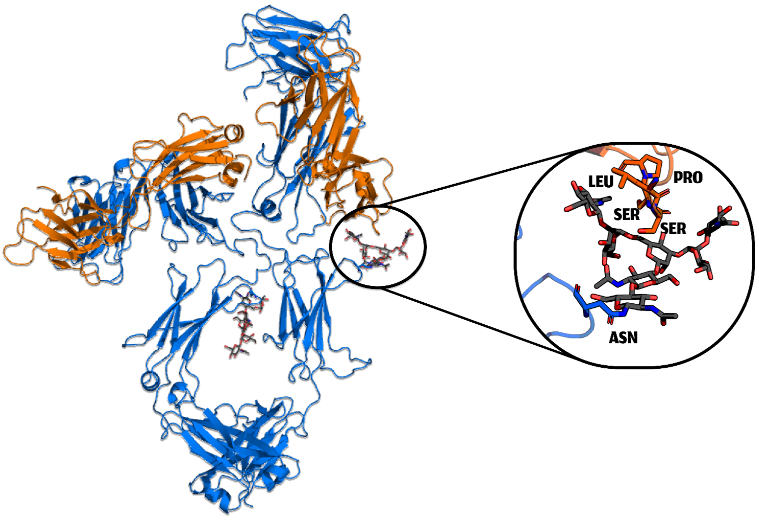


**B**

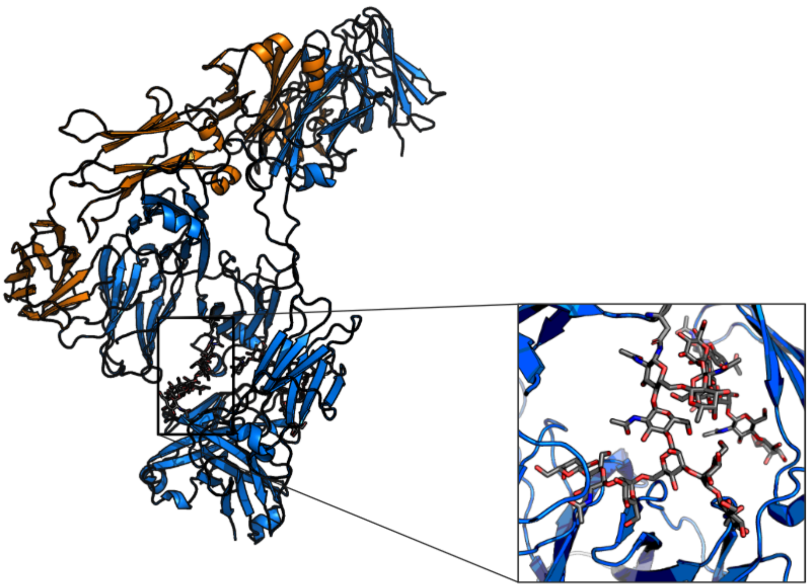

Figure S21 : Examples of conformations in which the Fabs come closer to the carbohydrates, for PMB (A) and Mab231 (B). Heavy chains are blue and light chains orange; the inset zooms on the glycan and the neighbouring residues (labelled in A: LEU, PRO, SER, SER, ASN) shown as sticks.

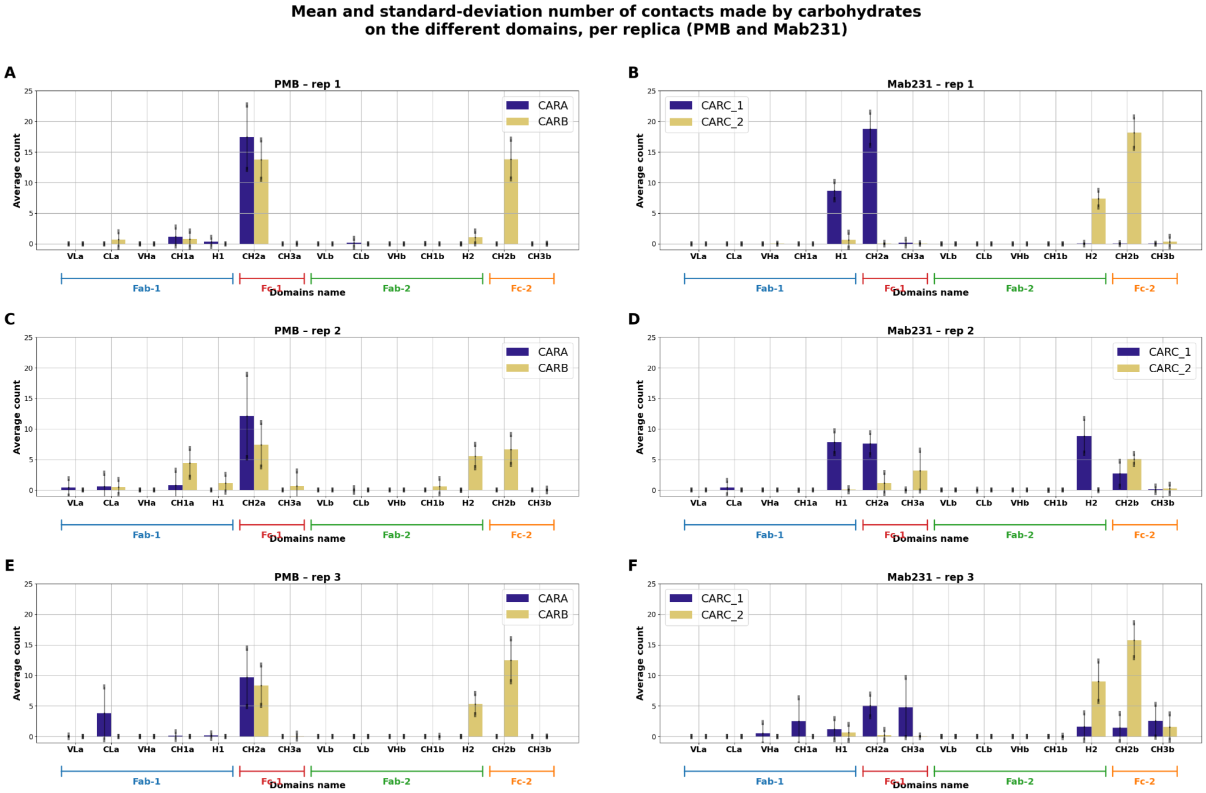


Figure S22 : Mean and standard-deviation number of contacts made by the carbohydrates on the different domains, per replica, for PMB (A, C, E) and Mab231 (B, D, F), corresponding to replicas 1, 2 and 3. For PMB, CARA is shown in blue and CARB in orange; for Mab231, CARC_1 in blue and CARC_2 in orange. Bars are averages over the trajectory and black lines are the standard-deviation values. Domain boundaries (Fab-1, Fc-1, Fab-2, Fc-2 and their subdomains) are indicated along the x-axis.

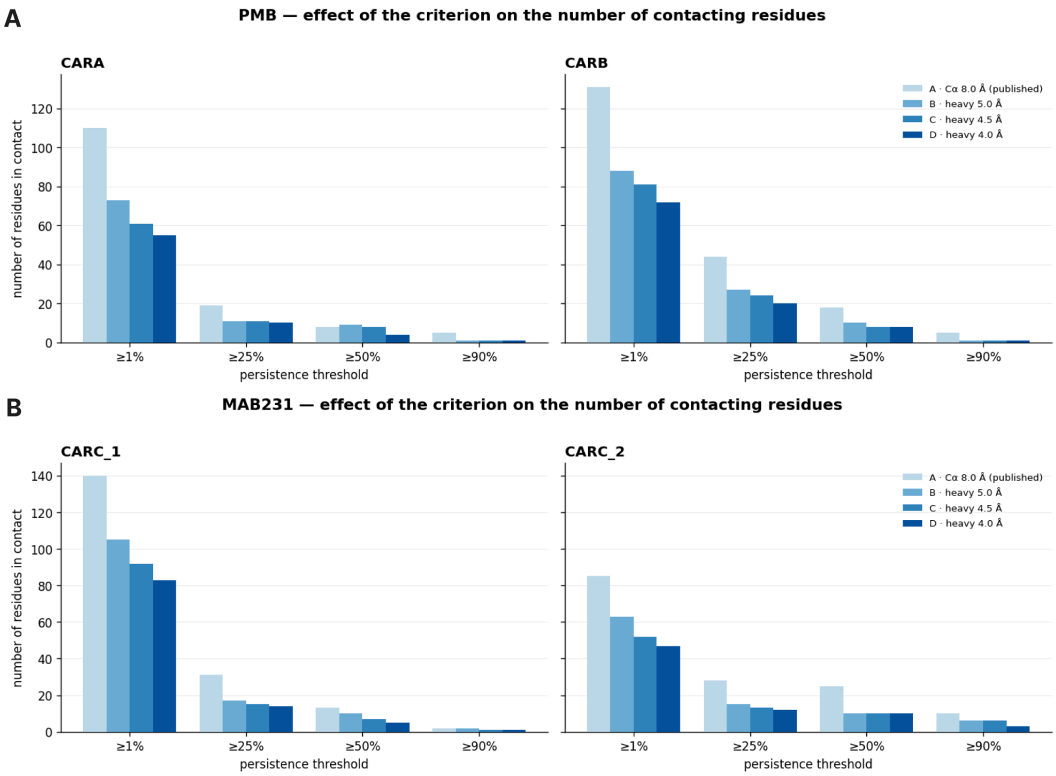


Figure S23 : Effect of the contact criterion on the number of residues in contact with each glycan, for PMB (A) and Mab231 (B). Bars give the number of residues above the 1 %, 25 %, 50 % and 90 % persistence thresholds, under criterion A (Cα within 8 Å, as in the main analysis) and under heavy-atom criteria at 5.0, 4.5 and 4.0 Å. Data are pooled across replicates, with the first 200 ns discarded.

*CH2 opening or closing analysis.*

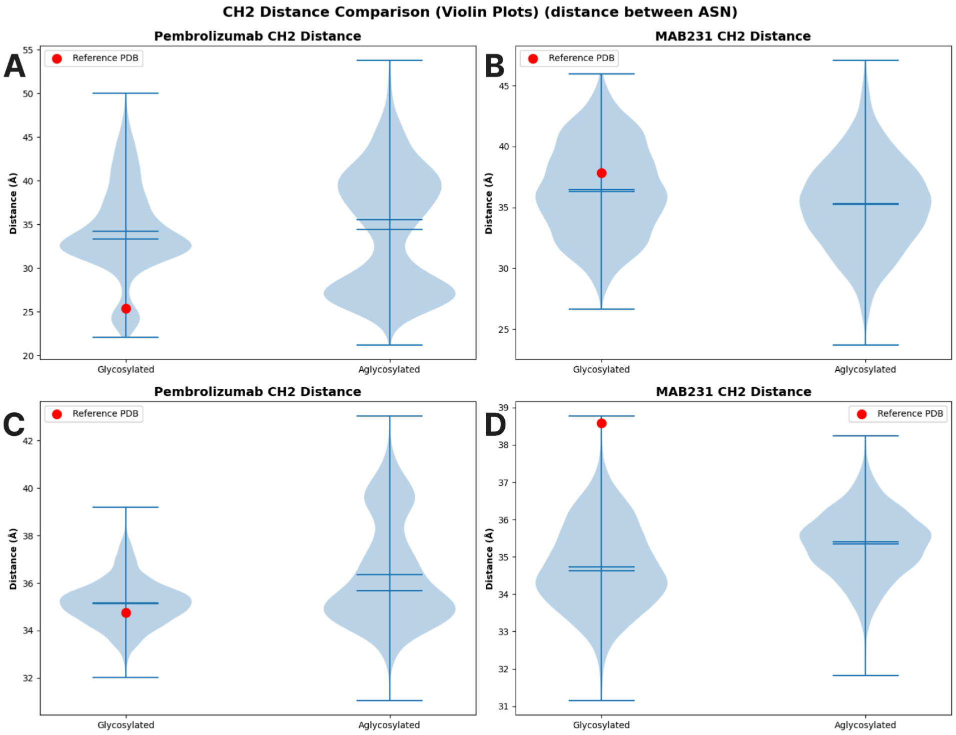


Figure S24 : Violin plots of the distribution of the distances used to characterise Fc opening, for the glycosylated and aglycosylated states. Top: distance between the Asn residues bearing the glycosylation, for PMB (A) and Mab231 (B). Bottom: distance between the centres of mass of the two CH2 domains, for PMB (C) and Mab231 (D). The red dot indicates the reference value in the crystallographic (PDB) structure. Horizontal lines within each violin mark the median and quartiles.

| Column | Test | Statistic | p-value | median r | median OVL |
|---|---|---|---|---|---|
| **θ Fab-1** | Mann-Whitney U | 1.03e09 | 1e-300 | -0.649 | 0.427 |
| **θ Fab-2** | Mann-Whitney U | 1.902e09 | 1e-300 | -0.406 | 0.596 |
| **θ inter Fab** | Mann-Whitney U | 2.537e09 | 1e-300 | -0.139 | 0.566 |

Table S3 : PMB summary statistics and p-values for the angles adopted by the Fabs (θFab-1, θFab-2, θInterFab), using the Mann–Whitney U test. The reported statistic, p-value, median rank-biserial effect size (median r) and median overlap coefficient (median OVL) are the medians of the three per-replicate values.

| Column | Test | Statistic | p-value | median r | median OVL |
|---|---|---|---|---|---|
| **θ Fab-1** | Mann-Whitney U | 1.973e09 | 1e-300 | -0.38 | 0.578 |
| **θ Fab-2** | Mann-Whitney U | 4.308e09 | 1e-300 | 0.346 | 0.401 |
| **θ inter Fab** | Mann-Whitney U | 3.607e09 | 1e-300 | 0.134 | 0.516 |

Table S4 : Mab231 summary statistics and p-values for the angles adopted by the Fabs (θFab-1, θFab-2, θInter-Fabs), using the Mann–Whitney U test. The reported statistic, p-value, median r and median OVL are the medians of the three per-replicate values.

***Supplementary References.***


1. Abraham, M. J. *et al.* GROMACS: High performance molecular simulations through multi-level parallelism from laptops to supercomputers. *SoftwareX* **1–2**, 19–25 (2015).
2. Grant, B. J., Rodrigues, A. P. C., ElSawy, K. M., McCammon, J. A. & Caves, L. S. D. Bio3d: an R package for the comparative analysis of protein structures. *Bioinformatics* **22**, 2695–2696 (2006).
3. Saporiti, S. *et al.* IgG1 conformational behavior: elucidation of the N-glycosylation role via molecular dynamics. *Biophys. J.* **120**, 5355–5370 (2021).
4. Grossfield, A. & Zuckerman, D. M. Chapter 2 Quantifying Uncertainty and Sampling Quality in Biomolecular Simulations. in *Annual Reports in Computational Chemistry* vol. 5 23–48 (Elsevier, 2009).
